# Stochastic sampling provides a unifying account of visual working memory limits

**DOI:** 10.1101/771071

**Authors:** Sebastian Schneegans, Robert Taylor, Paul M Bays

## Abstract

Research into human working memory limits has been shaped by the competition between different formal models, with a central point of contention being whether internal representations are continuous or discrete. Here we describe a sampling approach derived from principles of neural coding as a new framework to understand working memory limits. Reconceptualizing existing models in these terms reveals strong commonalities between seemingly opposing accounts, but also allows us to identify specific points of difference. We show that the discrete versus continuous nature of sampling is not critical to model fits, but that instead random variability in sample counts is the key to reproducing human performance in both single- and whole-report tasks. A probabilistic limit on the number of items successfully retrieved is an emergent property of stochastic sampling, requiring no explicit mechanism to enforce it. These findings resolve discrepancies between previous accounts and establish a unified computational framework for working memory that is compatible with neural principles.

---

Working memory refers to the nervous system’s ability to form stable internal representations that can be actively manipulated in the pursuit of behavioral goals. A classical view of visual working memory (VWM) held that it was organized into a limited number of memory slots, each capable of holding a single object [1, 2]. This model was subsequently modified to allow multiple slots to hold the same object and be combined on retrieval to achieve higher precision [3]. This “slots+averaging” model incorporated aspects of an alternative view, which holds that VWM is better conceptualized as a continuous resource that can be flexibly distributed between different objects or visual elements [4, 5], accounting for set size effects in delayed reproduction tasks [6] (Fig. 1A) and flexibility in prioritizing representations [7]. Variable precision models [8, 9] additionally proposed that the amount of memory resource is not fixed but varies randomly from item to item and trial to trial. An alternative approach [10] sought to explain VWM errors from neural principles as decoding variability in population representations [11], with the limited memory resource equated to the total neural activity dedicated to storage. Here we show that each of these influential accounts of VWM can be interpreted within a common framework based on the statistical principle of sampling [12–18].

**Figure 1:**
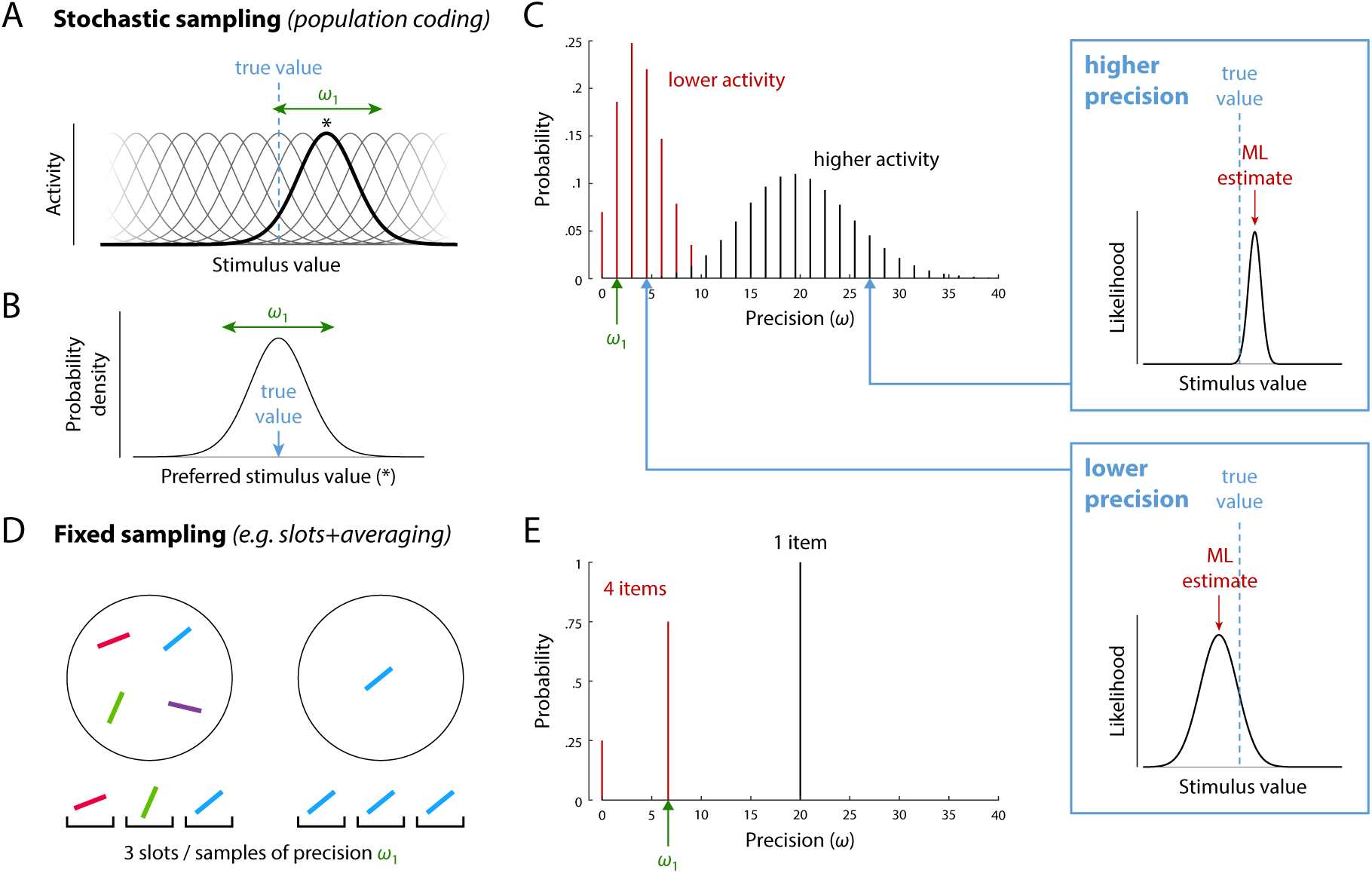
Sampling interpretation of working memory models. (A–C) A theoretical account of neural population coding can be re-interpreted as sampling. (A) The stimulus-evoked response of spiking neurons in an idealized population depends on their individual tuning (one neuron’s tuning function and preferred value [*] is highlighted). (B) Probability distribution over stimulus space obtained by associating a spike with the preferred stimulus of the neuron that generated it. (C) Precision of maximum likelihood estimates based on spikes emitted in a fixed decoding window. Precision, defined as the width of the likelihood function (insets), is discretely distributed as a product of the tuning precision (*ω*_1_) and the number of spikes, which varies stochastically. Assuming normalization of total activity encoding multiple items, larger set sizes correspond to less mean activity per item. (D–E) An account based on averaging limited memory slots can also be described as sampling. (D) Illustrates allocating a fixed number of samples or slots (here, three) to memory displays of different sizes. (E) Precision is discretely distributed as a product of the tuning width, *ω*_1_, and the number of samples allocated per item.

## Sampling interpretation of population coding

We first show how a population coding model [10] can, with some simplifying assumptions, be reinterpreted in terms of sampling (Fig. 1A–C). We consider a mathematically idealized population of independent neurons encoding a one-dimensional stimulus feature *θ*, where the amplitude of each cell’s activity is determined by its individual tuning function. Neurons are assumed to share the same tuning function, merely shifted so the peak lies at each neuron’s preferred feature value *φ*_*i*_:

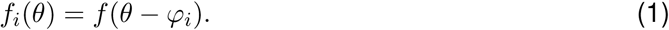

Discrete spikes are generated from the cells’ activity via independent Poisson processes. If we pick at random any spike generated by the neural population in response to a stimulus value *θ*, we can determine the probability that it was produced by a neuron with preferred feature value *φ*. If we assume dense uniform coverage of the underlying feature space by neural tuning curves, this yields a continuous probability distribution *p*(*φ*) over the space of preferred feature values (Fig. 1C). This distribution has the same shape as the neural tuning curves and is centered on the true stimulus value:

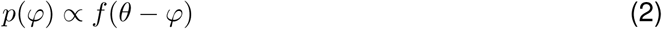

Thus, if we associate each spike with the preferred feature value of the neuron that generated it (the principle of population vector decoding; [19]), we can interpret the spiking activity of the population as a set of noisy samples of the true stimulus value, drawn from the distribution *p*(*φ*).

Retrieval of a feature value is modeled as decoding of the spikes generated within a fixed time window. In the idealized case with Gaussian tuning functions, the maximum likelihood (ML) decoder generates an estimate by simple averaging of the spike values:

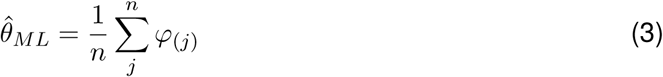

where *φ*(*j*) is the preferred feature value of the neuron that generated the *j*th spike.

Due to the superposition property of Poisson processes, the number of spikes – or samples – generated by the neural population within the decoding window is also a Poisson random variable. If the total spike rate in the neural population is normalized [20], or fixed at a population level *γ*, it implements a form of limited resource [10]. This resource is continuous – unlike the discrete number of samples – and can be distributed between memory items depending on task demands (e.g., prioritizing one item that is cued as a likely target). We will focus on the simplest case, in which the total spike rate is distributed evenly among all memory items, resulting in a mean number of samples available for decoding each stimulus that is inverse to the set size *N*. This has been shown to quantitatively capture the set size effect in single-report delayed reproduction tasks (Fig. 2A).

**Figure 2:**
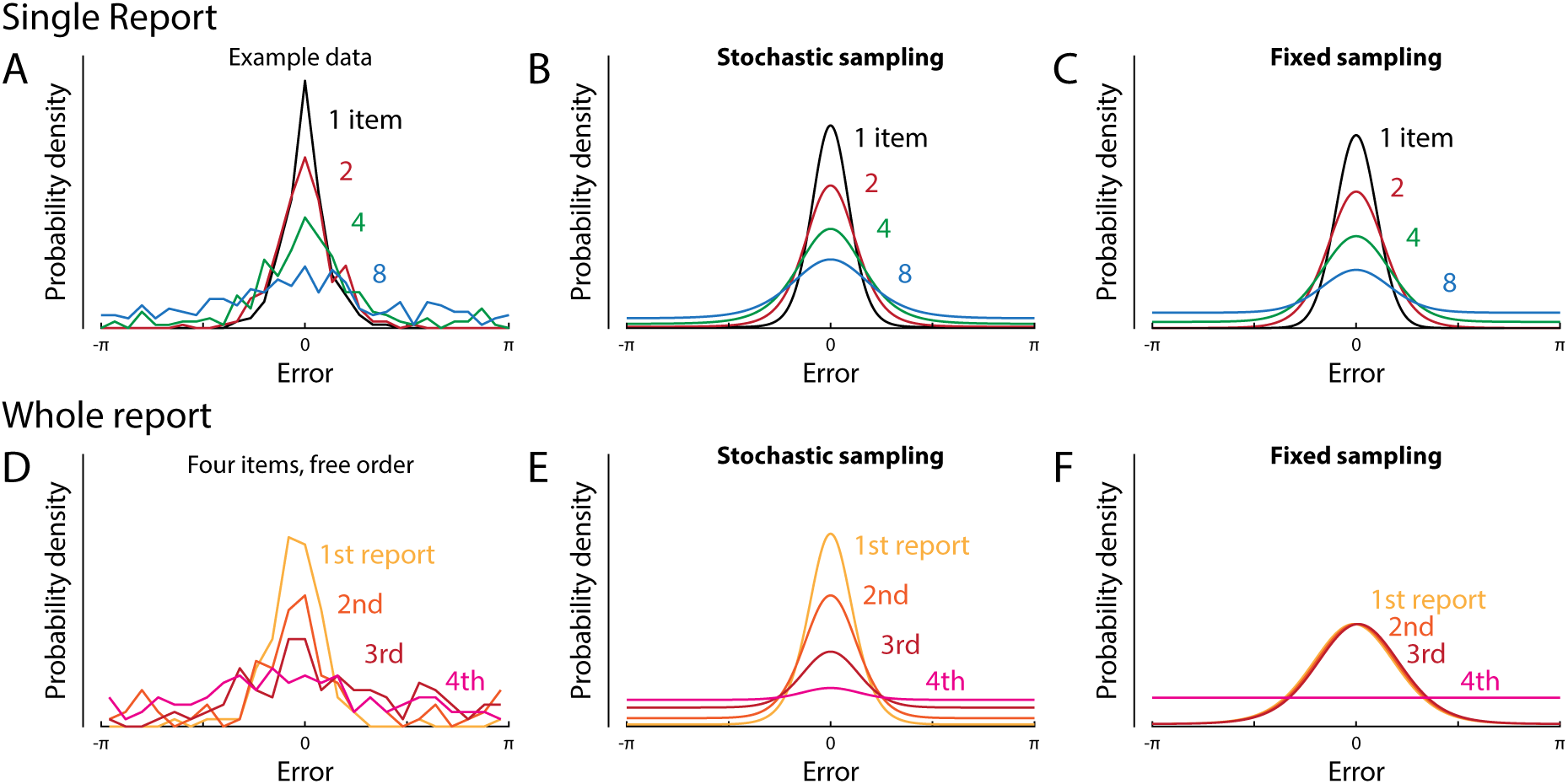
Response distributions and model fits in delayed reproduction tasks. (A) Distributions of response errors in a single-report task for a representative participant at different set sizes [10]. (B, C) ML fits of the data in (A) with the stochastic sampling model and fixed sampling model, respectively. (D) Distributions of response errors in a whole-report task for a representative participant at set size 4, showing how errors increase with the (freely-chosen) order of sequential report [24]. (E, F) ML fits of the participant’s data with the stochastic sampling model and fixed sampling model, respectively. Fits are based on results from all set sizes, not only the single set size shown in (D).

The actual number of samples available in this model for decoding each item in a single trial, *n*_*k*_, is a discrete random variable independently drawn from a Poisson distribution, with its mean determined by the spike rate for that item:

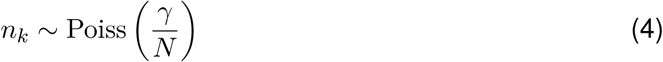

The neural population model can therefore be interpreted as a *stochastic* sampling model.

## Fixed sampling models

The most prominent discrete-representation account of VWM, the slots+averaging model [3], can also readily be interpreted in terms of sampling (Fig. 1D–E). Each slot is postulated to hold a representation of a single item with a fixed precision, and so provides a noisy sample of the item’s feature value (or values; the sampling interpretation is agnostic as to feature-vs object-based views of VWM; [21, 22]). Multiple slots, or samples, that correspond to the same object are averaged at retrieval to enhance the precision of the estimated stimulus feature. Thus, the format of representation and the decoding mechanism are identical to the stochastic sampling model. There is one critical difference, however: The slots+averaging model assumes that the total number of samples available for representing multiple items is fixed, i.e.

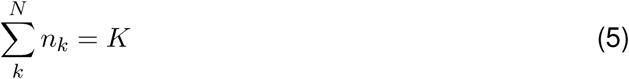

This has also been the most common assumption in previous sampling-based models in the attentional and memory literature ([12–14]; but see [23]). We will refer to this as a *fixed* sampling model.

## Predictions for precision and error distributions

We now consider the distribution of representational precision in these models. For any particular set of samples, **φ**, the information they provide about the stimulus is described by the likelihood function, *ℒ* (*θ*; **φ**) = *p*_*θ*_(**φ** |*θ*), equivalent to the conditional probability of obtaining those samples given different values of the stimulus. The width of the likelihood function is a measure of uncertainty in the estimate: a set of samples with a broad likelihood function (Fig. 1C, bottom inset) is compatible with many different feature values, whereas a narrow likelihood function (top inset) identifies a value more precisely. While a pattern of samples may have a sharp likelihood function with a peak far from the true estimate (a kind of “false alarm”), statistically this is unlikely.

If the sample values follow a normal distribution with variance *σ*^2^ centered on the true stimulus value, then the likelihood function is also normal, with a width that depends only on the number of samples available for decoding,

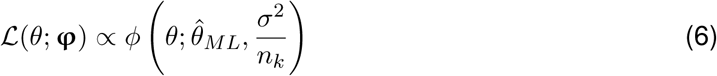

Furthermore, for a specified number of samples, the ML estimate is distributed around the true stimulus value as a normal with the same width as the likelihood,

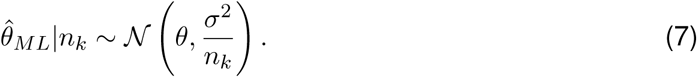

This correspondence between uncertainty, as expressed in the likelihood width, and trial-to-trial variability is not universal, but does apply to all the models considered in this study, and justifies defining the precision of an individual estimate (which we will denote *ω*) as the precision of its corresponding likelihood function (see Fig S1 for a detailed illustration). Adopting this definition explicitly (see also [25, 26]) allows us to treat precision as a random variable with a defined probability function, describing variation in the reliability of estimates while also predicting the distribution of errors across trials. This will prove critical in fitting data from whole-report tasks (Fig. 2D and below).

For the stochastic sampling model based on population coding, likelihood precision has a Poisson distribution (Fig. 1C), scaled by the precision of a single sample which is determined by the neural tuning function, *ω*_1_ = 1*/s*^2^,

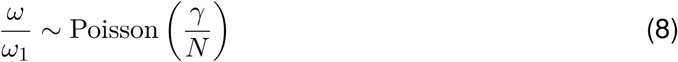

Example distributions of decoding error are shown in Fig. 2B & E, where we have made a transition from 1-D Euclidean to a circular stimulus space, corresponding more closely to the feature dimensions (e.g. orientation, hue) commonly used experimentally. The distribution of errors can be described as a scale mixture of normal distributions with precision proportional to the sample count (Fig. S1; due to the circular stimulus space, this is a close approximation rather than exact; see SI). The dispersion of errors increases with decreasing activity (e.g. as a result of increasing set size; Fig. 2B) and the distribution deviates from normality, with this effect being particularly evident at lower activity levels (blue curve) where long tails are observed.

For the fixed sampling model, making the common assumption that samples are distributed as evenly as possible among items [9, 27], we obtain a discrete distribution over at most two precision values (Fig. 1E), which are multiples of the precision of one sample, *ω*_1_. As in the stochastic model, mean precision is inversely proportional to set size, but because the distributions over precision differ, the fixed and stochastic models make distinct, testable predictions for error distributions (Fig. 2).

## Response errors discriminate between models

We tested the ability of stochastic and fixed sampling models to capture response errors in delayed reproduction tasks (Fig. S2). We fit the models to a large dataset of single-report tasks originating from different labs (Table S1) and also to a set of whole-report experiments [24], in which participants reported the feature values of all items in a sample array, either in a prescribed random order or in an order freely chosen by each participant on each trial. While only a single study, the whole-report results include information regarding correlations in errors between items represented simultaneously in VWM that could differentiate the models. On free choice trials, we assumed that participants gave their responses in order of decreasing precision (corresponding to decreasing number of samples and increasing likelihood width). This assumption is supported by previous findings that humans have knowledge about the uncertainty with which individual items are recalled [8, 25].

Overall, the stochastic model fit data substantially better than the fixed sampling model for both single-report (Fig. 3A; difference in log likelihood per participant [ΔLL] = 16.3 ± 2.37 [M ± SE]) and whole-report tasks (Fig. 3B; ΔLL = 162 ± 13.6), indicating that stochasticity is critical for capturing behavioral performance (see also Fig. S3). The response error distributions in the whole report task with freely chosen response order have previously been argued to provide evidence for a fixed item limit [24], since they approach uniform distributions for the later responses at high set sizes (Fig. 2D; see Figs. S4–S5 for full behavioral results and model fits). However, this qualitative observation is also predicted by the stochastic sampling model with responses ordered by precision, as the lowest precision retrievals will be based on few, or no, samples (Fig. 2E). Quality of fits could be further improved by taking into account deterioration of recall precision with increasing retention intervals (Figs. S3J and S5), modeled as random drift of encoded feature values over time ([28]; see SI).

**Figure 3:**
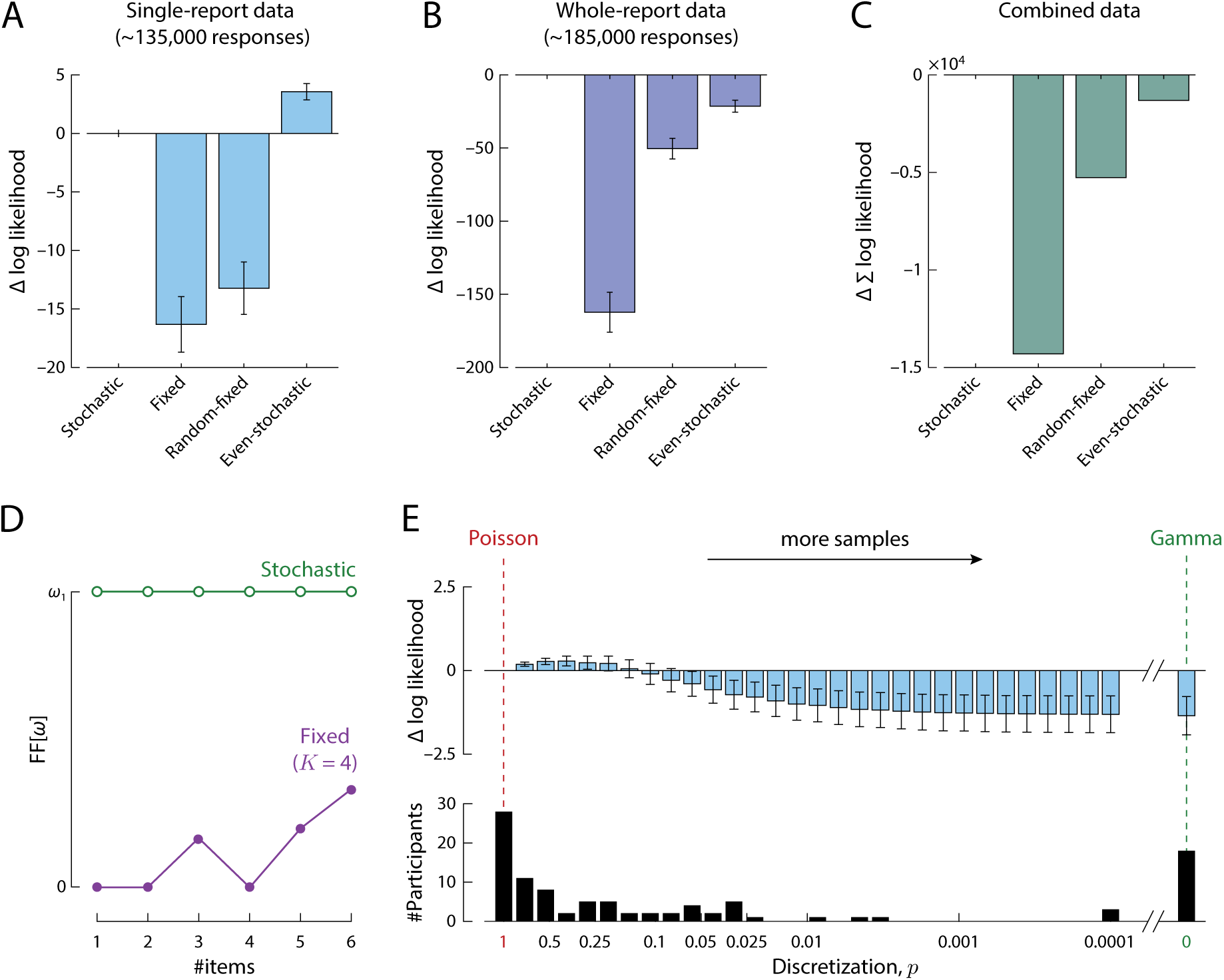
Model comparison based on single- and whole-report data. (A) Mean difference in log-likelihood of each model from the stochastic sampling model (with independence between items), for a benchmark data set of single-report experiments. More positive values indicate better fits to data. Errorbars indicate ±1 SE across participants. (B) The same comparison for a set of whole-report experiments. (C) Total difference in log-likelihood between models across single- and whole report experiments. (D) Fano factor (ratio of variance to mean) of precision distribution. A constant Fano factor is characteristic of the stochastic model and contrasts with the varying Fano factor (dependent on set size and number of samples) in fixed sampling. (E) Mean difference in log-likelihood for differing levels of discretization in the generalized stochastic model. Differences are plotted relative to the maximum discretization (*p* = 1; left) corresponding to the standard stochastic model with Poisson-distributed precision. Lower discretization (*p <* 1) corresponds to more samples each of lower precision, converging to a continuous Gamma distribution over precision as *p* approaches zero (right). All models have the same number of free parameters and include a fixed per-item probability of swap errors (see SI).

In contrast, the quantitative changes in error distribution with response order and set size were relatively poorly fit by the fixed sampling model (Fig. 2F). In particular, when the set size exceeds the fixed sample count, each item is represented by either one or zero samples, so this model cannot reproduce the graded decline in precision with response order that is also present in individual participants’ data (and does not merely arise at the group level due to averaging across participants with different capacities).

We tested two intermediate model versions in order to further dissociate the specific aspects in which the fixed and stochastic sampling models differ, and determine the significance of each for capturing human performance. In the *random–fixed* model, the total number of samples was fixed but distributed randomly between items. This model provided an improved fit to data compared to the fixed model with even allocation (moderately for single-report, ΔLL = 3.07 ± 1.10; strongly for whole-report, ΔLL = 112 ± 11.9), but was still substantially worse than the stochastic model in both cases (single-report, ΔLL = 13.2 ± 2.24; whole-report, ΔLL = 50.4 ± 7.03). In the *even–stochastic* model, the total number of samples was a Poisson random variable, but the samples were distributed as evenly as possible between items. This model achieved a better fit to single-report data than the stochastic model with independent sample counts for each item (ΔLL = 3.57 ± 0.697), but provided a much worse fit to whole-report data (ΔLL = 21.4 ± 4.12). Combining evidence across all participants and tasks, the stochastic model with independent sample counts was strongly preferred over this and the other alternative models (total ΔLL *>* 1450; Fig. 3C).

## Generalizing the stochastic model

For the models examined above, typical fitted parameters indicate that estimates are based on relatively small numbers of samples (e.g. mean of ∼13 samples based on fits to single-report data). One result is that the precision of decoded estimates could take on only a limited set of possible values, and error distributions reflect a discrete mixture of distributions with different widths. From a neural perspective, while consistent with the remarkable fidelity with which single neurons’ activity encodes visual stimuli [29, 30], such small sample counts nonetheless seem unlikely when interpreted as spike counts (see Towards biophysically realistic models, below). To investigate whether discreteness and/or low numbers of samples are important for reproducing human performance, we therefore implemented a generalization of the stochastic model in which the number of samples was free to vary.

The distribution over precision values in the generalized stochastic model was obtained as a scaling of the negative binomial distribution,

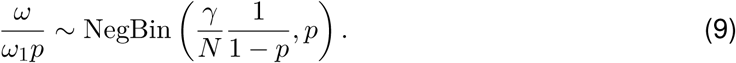

This distribution has previously been proposed to model neural spiking activity [31], and it retains the characteristic relationship between mean and variability in the scaled Poisson distribution: The Fano factor (the ratio of variance to mean) is constant, equal to the value of a single sample, *Var* [*ω*]*/E* [*ω*] = *ω*_1_. This distinguishes the stochastic models from the fixed sampling model, where the Fano factor is at or close to zero (mean ∼0.25 of *ω*_1_ based on ML parameters and typical set sizes) and varies in an idiosyncratic manner between set sizes, due to the varying combinatorial possibilities of allocating a fixed number of samples to a fixed number of items (Fig. 3D, purple).

The parameter *p* in the generalized stochastic model controls the discretization of the precision distribution: *p* = 1 corresponds to the Poisson model described above and illustrated in Fig. 4A (strictly Eq. 8 is the limit of Eq. 9 as *p* →1), while *p <* 1 corresponds to a stochastic model with a greater mean number of samples, 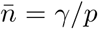, each with a lower individual precision, *ω*_1_*p*. The mean and variance in precision 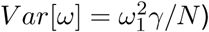, and thus also the Fano factor, are independent of the discretization *p*. Examples of precision distributions with different discretizations are shown in Fig. 4B & C.

**Figure 4:**
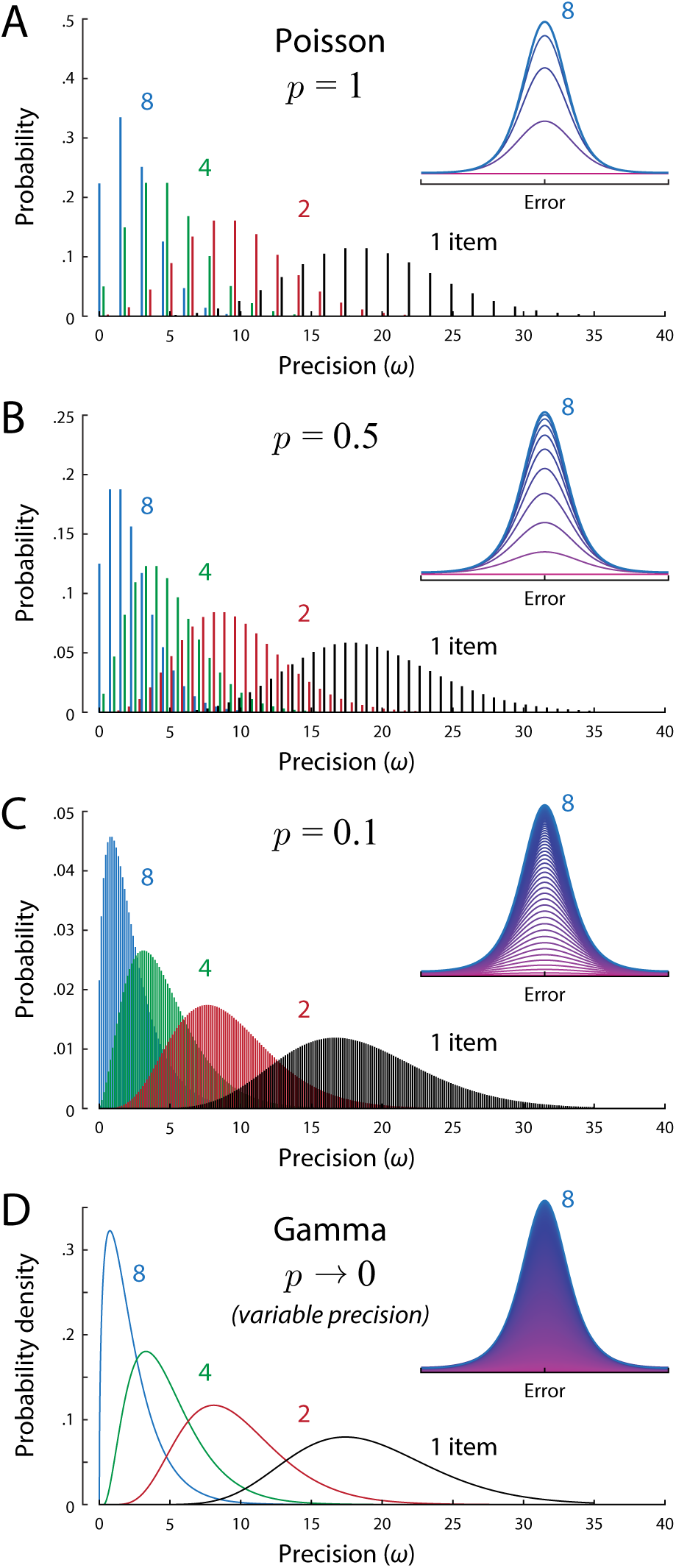
Precision distributions in the generalized stochastic model, for different levels of discretization, *p*, and different set sizes. Insets: Construction of the corresponding distributions of response error (for set size 8), with thin lines showing normal distributions with different precisions incrementally accumulated in ascending order (magenta to blue). (A) Example of discrete Poisson-distributed precision values (*p* = 1). For typical ML parameters, estimates are based on a small mean number of samples (here, *γ* = 12) each of moderate precision (*ω*_1_ = 1.5). (B & C) With decreasing discretization (*p <* 1), estimates are based on larger mean numbers of samples and discrete precision values are more finely spaced. (D) In the limit as discretization falls to zero, the mean number of samples becomes infinite and the distribution over precision approaches a continuous Gamma distribution. The ratio of variance to mean precision (Fano factor) is fixed (at *ω*_1_ = 1.5) across all set sizes and levels of discretization.

As the discretization parameter becomes very small (*p* → 0), the number of samples becomes very large and the distribution of precision described by Eq. 9 approaches a continuous function (Fig. 4D; see SI), specifically the Gamma distribution,

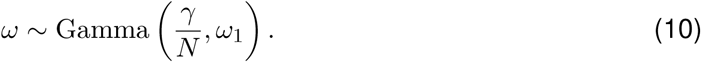

Two previous studies [8, 9] independently proposed that a continuous scale mixture of normal distributions with Gamma-distributed precision provided a good account of VWM data, but did not provide a theoretical motivation for this choice of distribution. In particular [9] proposed distributing precision as 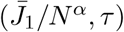, with 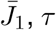, *τ* and *α* free parameters. With *α* = 1 this is identical to Eq. 10 (see SI for results regarding this parameter). We can now explain Gammadistributed precision as a limit case of the stochastic sampling model with large numbers of low-precision samples.

Fig. 3E (top) shows the results of fitting the generalized stochastic model with different levels of discretization, *p*, to the single-report dataset. The best fit was obtained with a discretization roughly one third that of the Poisson model, *p* = 0.39. However, varying discretization produced differences in fit an order of magnitude smaller than those between fixed and stochastic sampling (varying by ∼1.5 versus ∼15 log likelihood points). Fitting the same model with *p* as a free parameter that could vary between participants, we found that ML estimates of discretization were very broadly distributed (Fig. 3E, bottom), with a majority of participants (72%) best described by a sampling model with less discreteness than the Poisson, and a minority (18%) better captured by the continuous limit (*p* → 0) than any discrete value of *p* we tested (as low as 0.0001, corresponding to ∼100,000 samples). Formal model comparison was equivocal with respect to an advantage of including the discretization parameter in comparison to either the Poisson model (i.e. *p* = 1; ΔAIC = –0.61 ± 0.49; ΔBIC = +4.2 ± 0.46; negative values favor the added parameter) or the continuous Gamma model (i.e. *p* → 0; ΔAIC = –3.3 ± 0.93; ΔBIC = +1.5 ± 0.89). Overall, these results do not allow strong conclusions to be drawn regarding the discreteness of sampling, which has relatively little effect on error distributions (Fig 4 insets) or the quality of fits.

## Probabilistic item limits

In the fixed sampling model, at higher set sizes, a meaningful proportion of estimates are random “guesses” based on no samples (Fig. 5A & B). Specifically, if an estimate was generated for every item in the memory array, then as set size *N* increased, the number of estimates based on at least one sample, *S*_*ω>*0_, would reach a maximum at the fixed total number of samples,

**Figure 5:**
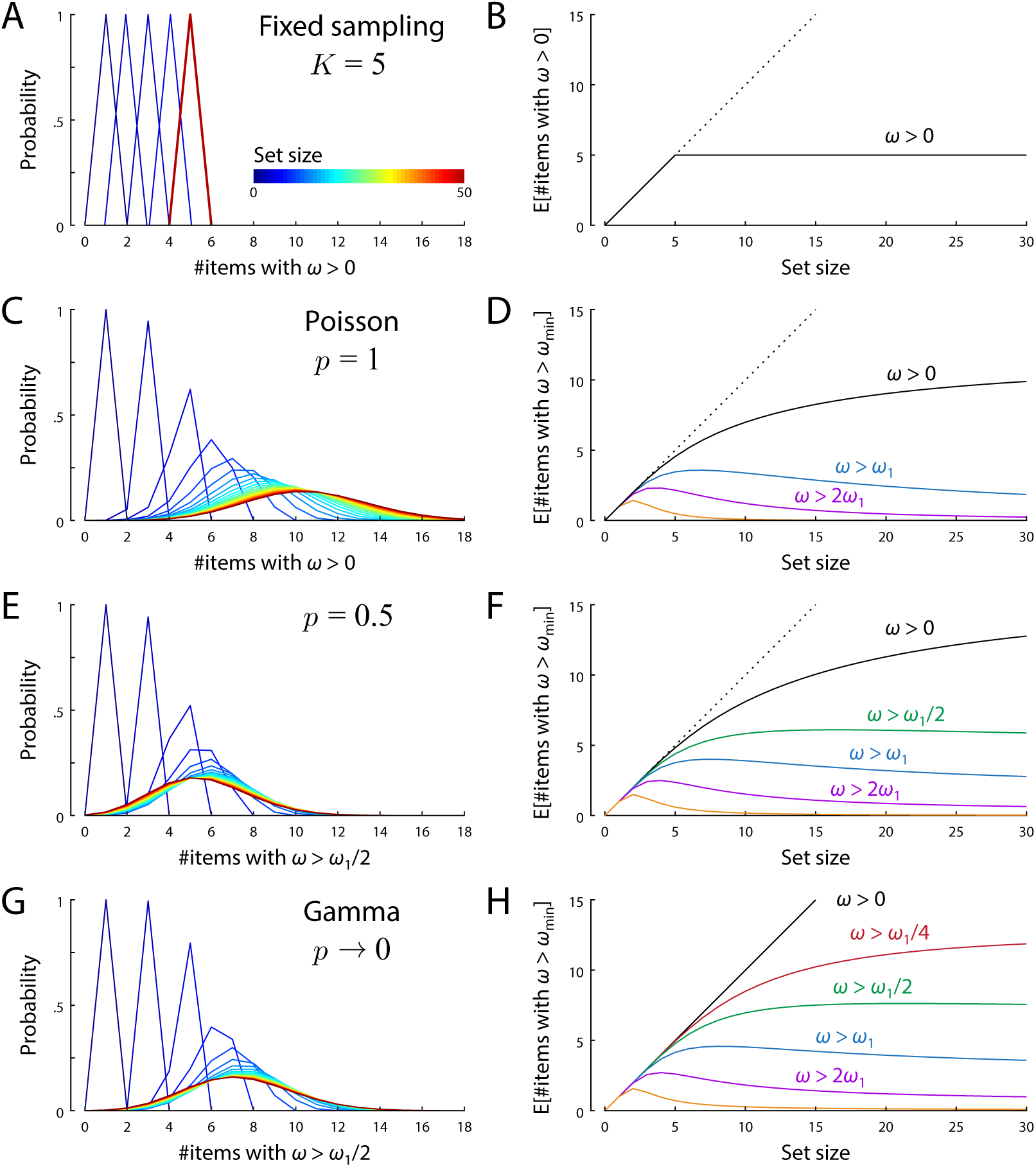
Item limits in sampling models. The left panel for each model shows how the probability distribution of the number of items recovered with greater than zero precision (A & C; greater than a fixed threshold for E & G) changes with set size (color coded, increasing blue to red; discrete probability distributions are depicted as line plots for better visualization). The right panels plot the mean number of items with above-threshold precision as a function of set size for different threshold values. Thresholds are defined as a proportion of the base precision *ω*_1_. (A, B) In the fixed sampling model the number of items with non-zero precision increases with set size, then plateaus when the number of items equals the number of samples. (C, D) The stochastic sampling model with Poisson variability also has a limit on the number of items with non-zero precision, although this limit is probabilistic and emerges asymptotically (converging to the distribution shown by the red curve in C for large set sizes, corresponding to the mean number of items plotted as black curve in D). (E–H) Stochastic models with lower discretization (E & F) or continuous precision (G & H), display similar probabilistic item limits for precision exceeding a fixed threshold, but with the expected number of items saturating at different values depending on threshold (colors in right-hand plots).

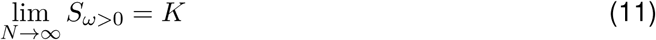

This is a trivial consequence of sharing out a fixed number of samples evenly between items.

In the stochastic model with Poisson variability (*p* = 1), the number of samples available for each item varies probabilistically and independently of the other items. There is again a probability of making an estimate based on zero samples,

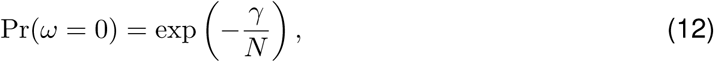

and the number of non-random retrievals in a set of *N* items has a Binomial distribution,

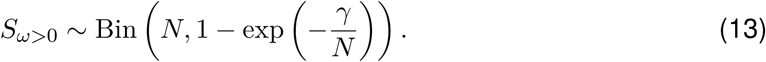

As set size increases, the mean number of estimates based on at least one sample reaches a maximum at the expected total number of samples (Fig. 5C & D). However, unlike the fixed sampling model, this limit is probabilistic and (as illustrated in Fig. 5C) the actual number will vary from one set of memory items to the next, converging to a Poisson distribution for large *N*,

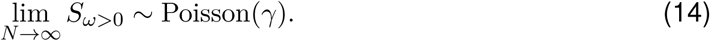

As we increase the expected number of samples by reducing the discretization, *p*, the probability of zero samples falls to zero: Pr(*ω* = 0) = *p*^*γ/*(*N*(1*–p*))^. However, if we choose a precision threshold that is less than or equal to the base precision *ω*_1_, it can be shown that the mean number of items with above-threshold precision converges to a finite positive number at large set sizes (see SI). This saturation is illustrated for different levels of discretization and various precision thresholds in Fig. 5E–H.

Item limits or “magic numbers” [2, 24, 32] are usually considered synonymous with slot-based accounts, occurring when some items must go unrepresented because other items have filled the available capacity. The present results show that a probabilistic item limit, i.e. an upper limit on the *average* number of items successfully retrieved that is not exceeded at any set size, can arise even when the probability of success for one item is independent of each other item. This holds true in the Poisson sampling model if we define success as obtaining one or more samples, but also more generally, even in a continuous model, if we define success as exceeding a threshold level of precision in estimation. Note however that the item limit does not in general have a simple relationship to the underlying number of samples, e.g. the probabilistic limit at *∼*5 items in Fig 5E is obtained from a model with a mean of 24 samples.

## Towards biophysically realistic models

The idealized description of population coding on which we based the stochastic sampling model overlooks a number of important considerations in order to reveal relationships between cognitive and neural-level accounts of VWM. For instance, the statistics of spike counts in the neural system often deviate from the Poisson distribution assumed in the original population coding model in that they are “overdispersed” (i.e. Fano factor *>* 1). Such an overdispersion in the sample count also occurs in the generalized stochastic model as the discretization *p* decreases (in order for the Fano factor of the precision distribution to remain constant, the Fano factor of the sample count has to increase). Spike counts in visual cortical neurons typically show a Fano factor in the range 1.5–3 (e.g. [33]), corresponding in our model to discretization *p* in the range 0.33–0.75.

In real neural populations there is also considerable variability between individual cells’ tuning curves [34]. Due to this heterogeneity, neurons differ in the amount of information each spike provides about a stimulus. From a sampling perspective, this means that estimates are based on samples that vary in precision, and this has the effect of “smoothing out” the discrete distribution of precision values predicted by the stochastic model (Fig. S6). This has similar consequences for estimation error as decreasing *p* in the generalized model. We fit the single report data with a variant of the population model with random variability in the neurons’ tuning curves (affecting baseline activity, gain, and tuning curve width, as well as adding heterogeneity in the coverage of the feature space by neural tuning curves; see SI), scaled by a global heterogeneity parameter *ν*. Incorporating biologically realistic heterogeneity into the population model improved fits to data (ΔAIC = 8.3 ± 1.8, ΔBIC = 3.4 ± 1.7 compared to the stochastic sampling model). The mean heterogeneity parameter in the ML fits was *ν* =0.66 ± 0.08, where *ν* = 0 means no heterogeneity, and *ν* = 1 was approximately matched to heterogeneity of orientation-selective neurons in recordings from V1 [34].

Finally, spikes in real neural populations are not independent events as assumed by the sampling interpretation, but rather correlated within and between neurons. This will tend to result in deviations from the simple additivity assumed by sampling. An implementation of short-range pair-wise correlations in the heterogeneous population model (see SI for details) greatly increased the numbers of decoded spikes required to reproduce behavioral data (on average, 168 times higher), without changing quality of fit (ΔAIC/BIC = 0.045 ± 0.28). We note however that the exact consequences of spike correlations for decoding depend on details of correlation structure that are difficult to measure experimentally [35–37], and suboptimal inference (in the form of a mismatched decoder) may play an important part [38].

## Discussion

Taking as a starting point a mathematical idealization of the way neural populations encode information, we have shown that retrieval of a visual feature from working memory can be described as estimation based on a stochastically varying number of noisy samples. Two other influential models of VWM can be reconceptualized in the same framework: The slots+averaging model, because it modified the original slot model to allow multiple representations with independent noise, is directly equivalent to a sampling model with a fixed number of samples [13]. And the variable precision model [8, 9] constitutes the continuous limit of a sampling model as samples are made less precise and more numerous, while maintaining the fixed proportionality between the variance and mean of precision in the decoded estimate.

Formulating all three models in the same mathematical framework (a formal “unification”) allowed us to pinpoint specific differences between them. We determined the effect of these differences on the models’ ability to account for human behavior by fitting multiple variants of the sampling model to a large database of delayed reproduction tasks. We found that stochasticity both in the total number of samples and their distribution among items has a major impact on the quality of fit, with the best fits obtained if the number of samples is drawn randomly and independently for each item in each trial. Note that this form of stochasticity is poorly captured by the concept of memory “slots”, because of the implication that a slot occupied by one item leaves fewer slots available for other items—this would predict dependencies between items in whole report that were not supported experimentally.

On the other hand, contrary to the assumptions of continuous resource models, we did find limited support for *discreteness* of memory representations [3]. The fully continuous model with Gamma-distributed precision proposed in previous studies provided fits to data that were overall a little worse than the discrete Poisson model, in both single- and whole-report tasks (see SI). When we attempted to fit discretization as a free parameter, however, we found ML estimates varied widely between participants, and many were best fit by continuous or near-continuous versions of the generalized stochastic sampling model. So while discreteness in memory representations is plausible – even inevitable if based on discrete spiking activity – recall errors do not provide strong evidence for any one particular level of discreteness, or as a corollary, any particular mean number of samples.

Our findings further highlight the need to distinguish between two concepts that have previously been elided: Discreteness in representation and discreteness in allocation. In the stochastic sampling model, the resource underlying capacity limits in VWM is equated with the *mean* number of samples (or the mean spike rate in the population coding interpretation), which can be distributed among items in a continuous fashion, even though the consequent number of samples obtained by each item is a discrete integer. This view on memory resources was strongly motivated by studies showing that prioritized items can be represented more precisely in VWM, at the cost of decreased precision for other items [7, 39–42]. The stochastic sampling model can account for such findings through an uneven distribution of resources among memory items, corresponding to a higher mean number of samples for some items at the cost of a lower mean for other items. The actual number of samples available on an individual memory retrieval varies randomly about the item’s mean. In the neural population model, this mechanism has previously been shown to successfully reproduce data from tasks in which one item is cued as the likely target [10].

While the stochastic sampling model is based on a highly idealized implementation of population coding, it nevertheless provides a link to a concrete neural mechanism that could form the basis of VWM performance. We have shown that adapting this model to achieve a higher degree of biophysical realism – by introducing heterogeneity in neural tuning curves and correlated spiking activity – improved the quality of fit to behavioral data. It has recently been shown that more neurally realistic population coding models preserve the key characteristics of the idealized model, and that signatures of neural tuning may even be visible in behavioral data [43].

Our results also provide a link between models of working memory used in the psychological literature and more biophysically detailed neural models such as continuous attractor networks [44–46], whose greater complexity typically precludes quantitative fits to behavioral data. These models are likewise based on principles of population coding and emphasize the role of neural noise in explaining variability in working memory performance. They are capable of producing probabilistic item limits similar to those described here, but it remains unresolved how these models could account for the graded variations in recall fidelity that we have found to be essential for capturing human behavioral performance.

In keeping with most previous work on VWM limits, we have not here attempted to reproduce the variations in bias and precision that are observed for different feature values, exemplified by the finding that cardinal orientations can be reproduced with greater precision than obliques. However, previous work has shown that these effects can be simply and elegantly captured within the population coding framework via the principle of efficient coding [47–49]. The idea is that neural tuning functions are adapted to the stimulus statistics of the natural environment in such a way as to maximally convey information in that environment (effectively by distributing neural resources preferentially to the most frequently-occurring stimulus features). Although it should be possible to formulate this model as a modification of stochastic sampling, without reference to neural populations, it seems that the modifications required would not have a natural explanation within the sampling framework. These observations, and the results of incorporating heterogeneity described above, illustrate the value of connecting abstract cognitive models to neural theory.

We also did not address here the question how individual features of a visual stimulus are bound together, which forms another point of contention in the debate on the format of VWM representations. In the model fits, we allowed failures of binding memory in the form of swap errors to occur with a fixed rate, although taking into account similarity of items with respect to the cue feature is likely to improve model fits [50, 51]. While the discrete memory representation in slot models have traditionally been associated with a strongly object-based view [1], the sampling framework is agnostic to whether objects or features are the units of VWM storage. Both views are compatible with the population coding interpretation, depending on whether the neurons in question are sensitive to a single feature [52] or a conjunction of features [51, 53].

A recent proposal that VWM errors can be explained in terms of a perceptual rescaling of stimulus space can also be expressed in terms of population coding, with some minor differences from the version presented here (see [54], for details and discussion). In particular, the idea of retrieval based on normally distributed “memory-match” signals maps exactly onto an idealized population code with continuous-valued activity and constant Gaussian noise [51, 55]. This predicts a continuous distribution over precision, not dissimilar to the Gamma distribution. Continuousness in representation does not appear a necessary component of this account, however, and it should be possible to reformulate it with arbitrary levels of discreteness, as in our generalized stochastic model.

There are other models of working memory that address capacity limits without explicitly postulating a limited memory resource [56]. Some accounts stress the importance of memory decay over time, and active rehearsal to counteract this decay [57, 58]. These theories do not have a clear analogue in the sampling framework, although effects of retention time have been incorporated into the neural population model [28]. Other accounts have sought to explain capacity limits by interference between different memorized items [59]. While the sampling framework does not explicitly address interference, the effect of normalization could be described as a form of non-specific interference between items. A model of feature binding based on the neural population model shows some notable congruencies with an interference account of visual working memory, and both models make similar predictions regarding swap errors [50, 51]. Further research will be needed to determine the exact relationship between these models.

Taken together, our results reveal a surprising convergence between prominent models of VWM. Despite the fact that these competing models were independently motivated by different behavioral and neural findings, they can be expressed within the shared formal framework of sampling, which reveals specific distinguishing factors as well as shared general principles. This convergence gives cause for confidence that the stochastic sampling model captures key characteristics of VWM and will provide a solid foundation for future research.

## Material and methods

We fit computational models of VWM to behavioral data from a large dataset of delayed estimation experiments. The dataset included 15 individual single-report experiments (Table S1; see SI for inclusion criteria), as well as four whole-report experiments (Table S2). Each model defines a parameterized distribution of response probabilities given the true feature values of the target and non-target items in each trial (see SI). For fits to whole-report data, we determined the probabilities of obtaining the given combination of responses within a single trial, taking into account the correlations of recall precision between different items within a trial predicted by each model.

We obtained a ML fit of each subject’s data for each model. The stochastic sampling, fixed sampling, and continuous sampling (Gamma) model, as well as the fixed–random and stochastic– even variants, each have three free parameters (including one parameter for the probability of swap errors). We fit these to both the single-report and whole-report data using the NelderMead simplex algorithm (see SI for details). The generalized stochastic model and the neural population model with heterogeneous tuning curves have four free parameters each. For these models, we employed a grid search to obtain fits only of the single-report data (fitting them to whole-report data was not computationally feasible). We further evaluated model variants employing a more accurate method for ML decoding for circular feature spaces (rather than the Gaussian approximation used for fits reported in the main manuscript), models without swap errors, models with an additional free parameter for the power law in set size effects, and models with a temporal decay of memory precision over varying response delays in the whole-report experiments (see SI).

## Acknowledgments

We thank Ronald van den Berg, Wei Ji Ma, Máté Lengyel & Masud Husain for helpful conversations, Zakhar Kabluchko & Martin Bays for statistical advice, and all the researchers who publicly shared data that facilitated this study. We used resources provided by the Cambridge Service for Data Driven Discovery (CSD3) operated by the University of Cambridge Research Computing Service.

## Data Availability

Data and code associated with this article will be made available on publication at https://osf.io/buxp9/.

## References

[1] Steven J Luck and Edward K Vogel. “The capacity of visual working memory for features and conjunctions”. Nature 390.6657 (1997), pp. 279–281.

[2] Nelson Cowan. “The magical number 4 in short-term memory: A reconsideration of mental storage capacity”. Behavioral and Brain Sciences 24.1 (2001), pp. 87–114.

[3] Weiwei Zhang and Steven J. Luck. “Discrete fixed-resolution representations in visual working memory”. Nature 453.7192 (2008), pp. 233–235.

[4] Paul M Bays, Raquel FG Catalao, and Masud Husain. “The precision of visual working memory is set by allocation of a shared resource”. Journal of Vision 9.10 (2009), pp. 7–7.

[5] Wei Ji Ma, Masud Husain, and Paul M Bays. “Changing concepts of working memory”. Nature Neuroscience 17.3 (2014), p. 347.

[6] Patrick Wilken and Wei Ji Ma. “A detection theory account of change detection”. Journal of Vision 4.12 (2004), pp. 11–11.

[7] Paul M Bays and Masud Husain. “Dynamic shifts of limited working memory resources in human vision”. Science 321.5890 (2008), pp. 851–854.

[8] Daryl Fougnie, Jordan W Suchow, and George A Alvarez. “Variability in the quality of visual working memory”. Nature Communications 3 (2012), p. 1229.

[9] Ronald van den Berg et al. “Variability in encoding precision accounts for visual shortterm memory limitations”. Proceedings of the National Academy of Sciences 109.22 (2012), pp. 8780–8785.

[10] Paul M Bays. “Noise in neural populations accounts for errors in working memory”. Journal of Neuroscience 34.10 (2014), pp. 3632–3645.

[11] Alexandre Pouget, Peter Dayan, and Richard Zemel. “Information processing with population codes”. Nature Reviews Neuroscience 1.2 (2000), p. 125.

[12] Marilyn L. Shaw. “Identifying attentional and decision-making components in information processing”. Attention and Performance VIII 8 (1980), pp. 277–295.

[13] John Palmer. “Attentional limits on the perception and memory of visual information”. Journal of Experimental Psychology: Human Perception and Performance 16.2 (1990), p. 332.

[14] David K Sewell, Simon D Lilburn, and Philip L Smith. “An information capacity limitation of visual short-term memory”. Journal of Experimental Psychology: Human Perception and Performance 40.6 (2014), p. 2214.

[15] Anne-Marie Bonnel and Jeff Miller. “Attentional effects on concurrent psychophysical discriminations: Investigations of a sample-size model”. Perception & Psychophysics 55.2 (1994), pp. 162–179.

[16] Edward Vul et al. “One and Done? Optimal Decisions From Very Few Samples”. Cognitive Science 38.4 (2014), pp. 599–637.

[17] Gergő Orbán et al. “Neural Variability and Sampling-Based Probabilistic Representations in the Visual Cortex”. Neuron 92.2 (2016), pp. 530–543.

[18] Michael N. Shadlen and Daphna Shohamy. “Decision Making and Sequential Sampling from Memory”. en. Neuron 90.5 (2016), pp. 927–939.

[19] Apostolos P Georgopoulos, Andrew B Schwartz, and Ronald E Kettner. “Neuronal population coding of movement direction”. Science 233.4771 (1986), pp. 1416–1419.

[20] Matteo Carandini and David J. Heeger. “Normalization as a canonical neural computation”. Nature Reviews Neuroscience 13.1 (2012), pp. 51–62.

[21] George A Alvarez and Patrick Cavanagh. “The capacity of visual short-term memory is set both by visual information load and by number of objects”. Psychological Science 15.2 (2004), pp. 106–111.

[22] Klaus Oberauer and Simon Eichenberger. “Visual working memory declines when more features must be remembered for each object”. Memory & Cognition 41.8 (2013), pp. 1212–1227.

[23] Philip L Smith. “The Poisson shot noise model of visual short-term memory and choice response time: Normalized coding by neural population size”. Journal of Mathematical Psychology 66 (2015), pp. 41–52.

[24] Kirsten CS Adam, Edward K Vogel, and Edward Awh. “Clear evidence for item limits in visual working memory”. Cognitive Psychology 97 (2017), pp. 79–97.

[25] Ronald Van den Berg, Aspen H Yoo, and Wei Ji Ma. “Fechner’s law in metacognition: A quantitative model of visual working memory confidence.” Psychological Review 124.2 (2017), p. 197.

[26] Paul M Bays. “A signature of neural coding at human perceptual limits”. Journal of Vision 16.11 (2016), pp. 4–4.

[27] Ronald van den Berg, Edward Awh, and Wei Ji Ma. “Factorial comparison of working memory models.” Psychological Review 121.1 (2014), p. 124.

[28] Sebastian Schneegans and Paul M Bays. “Drift in neural population activity causes working memory to deteriorate over time”. Journal of Neuroscience (2018), pp. 3440–3417.

[29] Ehud Zohary, Michael N. Shadlen, and William T. Newsome. “Correlated neuronal discharge rate and its implications for psychophysical performance”. Nature 370.6485 (1994), pp. 140–143.

[30] K. H. Britten et al. “The analysis of visual motion: a comparison of neuronal and psychophysical performance”. Journal of Neuroscience 12.12 (1992), pp. 4745–4765.

[31] Robbe LT Goris, J Anthony Movshon, and Eero P Simoncelli. “Partitioning neuronal variability”. Nature neuroscience 17.6 (2014), p. 858.

[32] George A Miller. “The magical number seven, plus or minus two: Some limits on our capacity for processing information.” Psychological Review 101.2 (1994), p. 343.

[33] Rufin Vogels, Werner Spileers, and Guy A. Orban. “The Response Variability of Striate Cortical Neurons in the Behaving Monkey”. Experimental Brain Research 77.2 (1989), pp. 432–436.

[34] Alexander S Ecker et al. “Decorrelated neuronal firing in cortical microcircuits”. Science 327.5965 (2010), pp. 584–587.

[35] Bruno B Averbeck, Peter E Latham, and Alexandre Pouget. “Neural correlations, population coding and computation”. Nature Reviews Neuroscience 7.5 (2006), p. 358.

[36] Alexander S Ecker et al. “The effect of noise correlations in populations of diversely tuned neurons”. Journal of Neuroscience 31.40 (2011), pp. 14272–14283.

[37] Rubén Moreno-Bote et al. “Information-limiting correlations”. Nature Neuroscience 17.10 (2014), p. 1410.

[38] Jeffrey M Beck et al. “Not noisy, just wrong: the role of suboptimal inference in behavioral variability”. Neuron 74.1 (2012), pp. 30–39.

[39] Paul M Bays et al. “Temporal dynamics of encoding, storage, and reallocation of visual working memory”. Journal of Vision 11.10 (2011), pp. 6–6.

[40] Aspen H Yoo et al. “Strategic allocation of working memory resource”. Scientific Reports 8.1 (2018), p. 16162.

[41] Stephen M Emrich, Holly A Lockhart, and Naseem Al-Aidroos. “Attention mediates the flexible allocation of visual working memory resources.” Journal of Experimental Psychology: Human Perception and Performance 43.7 (2017), p. 1454.

[42] Zuzanna Klyszejko, Masih Rahmati, and Clayton E Curtis. “Attentional priority determines working memory precision”. Vision Research 105 (2014), pp. 70–76.

[43] Robert Taylor and Paul M. Bays. “Theory of neural coding predicts an upper bound on estimates of memory variability”. Psychological Review Advance Online Publication. http://dx.doi.org/10.1037/rev0000189 (2020).

[44] Jeffrey S Johnson et al. “A dynamic neural field model of visual working memory and change detection”. Psychological Science 20.5 (2009), pp. 568–577.

[45] Ziqiang Wei, Xiao-Jing Wang, and Da-Hui Wang. “From distributed resources to limited slots in multiple-item working memory: a spiking network model with normalization”. Journal of Neuroscience 32.33 (2012), pp. 11228–11240.

[46] Dominic Standage and Martin Paré. “Slot-like capacity and resource-like coding in a neural model of multiple-item working memory”. Journal of Neurophysiology 120.4 (2018), pp. 1945–1961.

[47] Robert Taylor and Paul M Bays. “Efficient coding in visual working memory accounts for stimulus-specific variations in recall”. Journal of Neuroscience 38.32 (2018), pp. 7132–7142.

[48] Xue-Xin Wei and Alan A Stocker. “A Bayesian observer model constrained by efficient coding can explain’anti-Bayesian’percepts”. Nature Neuroscience 18.10 (2015), p. 1509.

[49] Deep Ganguli and Eero P Simoncelli. “Efficient sensory encoding and Bayesian inference with heterogeneous neural populations”. Neural Computation 26.10 (2014), pp. 2103–2134.

[50] Klaus Oberauer and Hsuan-Yu Lin. “An interference model of visual working memory”. Psychological Review 124.1 (2017), p. 21.

[51] Sebastian Schneegans and Paul M Bays. “Neural architecture for feature binding in visual working memory”. Journal of Neuroscience 37.14 (2017), pp. 3913–3925.

[52] Flora Bouchacourt and Timothy J Buschman. “A flexible model of working memory”. Neuron 103.1 (2019), pp. 147–160.

[53] Loic Matthey, Paul M Bays, and Peter Dayan. “A probabilistic palimpsest model of visual short-term memory”. PLOS Computational Biology 11.1 (2015).

[54] Paul M Bays. “Correspondence between population coding and psychophysical scaling models of working memory”. BioRxiv (2019), p. 699884.

[55] Haim Sompolinsky et al. “Population coding in neuronal systems with correlated noise”. Physical Review E 64.5 (2001), p. 051904.

[56] John Jonides et al. “The mind and brain of short-term memory”. Annual Review of Psychology 59 (2008), pp. 193–224.

[57] Alan D Baddeley. Working memory. Oxford: Oxford University Press, 1986.

[58] Pierre Barrouillet and Valérie Camos. “As time goes by: Temporal constraints in working memory”. Current Directions in Psychological Science 21.6 (2012), pp. 413–419.

[59] Klaus Oberauer et al. “Modeling working memory: An interference model of complex span”. Psychonomic Bulletin & Review 19.5 (2012), pp. 779–819.

