## Supplementary Information for "Stochastic sampling provides a unifying account of visual working memory limits"

### 1 Behavioral data

To evaluate different models of visual working memory, we compared the quality of model fits for a large set of behavioral data from continuous report tasks. This dataset is compiled of 11 experiments with over 130,000 trials in total. We included available data from published delayed reproduction experiments (either single-report or whole-report tasks; Fig. S2) that have the following characteristics: They test recall performance for at least two different set sizes with a fixed delay duration, all items are presented simultaneously and are equally likely to be tested, and the reported feature is either color or orientation. The target item can be indicated either by a location cue or a categorical color cue. The dataset for single-report tasks (Table S1) is similar to the dataset used in a previous model comparison [1], but we excluded experiments from two studies that have in the meantime been retracted, and added several more recent studies. We also fit behavioral data from four whole-report tasks (Table S2), in which participants had to report the feature values of all items presented in the sample array, with the order of responses either freely chosen by the participant, or determined randomly by the experiment software [2]. In the whole-report tasks, items were always cued or selected via their location.

### 2 Models

#### 2.1 General assumptions and notations

For a single trial in a continuous report task, we denote the set size of the memory sample array with $N$, and the feature values of the sample items with $\boldsymbol{\theta} = (\theta_1, \ldots, \theta_N)$. For classical continuous report tasks with a single report, we denote the reported feature value with $\hat{\theta}$. For whole-report tasks, the sequence of reported feature values is $\hat{\boldsymbol{\theta}} = (\hat{\theta}_1, \ldots, \hat{\theta}_N)$.

Each model defines a response probability distribution $p(\hat{\theta}|\theta)$ assuming that the response is generated based on a sample item with true feature value $\theta$. For all model fits, we incorporate swap errors, which we found to consistently improve the quality of fits (see Section Excluding swap errors). We assume that response distributions around the selected feature value are identical for target and non-target features, and that the probability of reporting a non-target feature increases linearly with set size, with each non-target item having an equal probability $p_{\mathrm{NT}}$ of being used as the basis for response generation (these assumptions are of course not tenable for very large $N$, but suffices for studies in our dataset, with $N \leq 8$). For the response $\hat{\theta}_i$ corresponding to a cued item $\theta_i$, we then obtain the probability distribution

$$p(\hat{\theta}_i|\boldsymbol{\theta}) = p_{\mathrm{T}}p(\hat{\theta}_i|\theta_i) + p_{\mathrm{NT}} \sum_{j\in\{1,\ldots,N\},j\neq i} p(\hat{\theta}_i|\theta_j), \tag{1}$$

where $p_{\mathrm{T}} = 1 - (N-1)p_{\mathrm{NT}}$ is the probability that the response is based on the feature value of the target item.

In the whole-report tasks with freely chosen response order, we assume for all models that responses are ordered by precision (either expressed as the number of samples assigned to them or as a continuous precision value), starting with the highest precision item. We further assume that this ordering is still maintained if a swap error occurs. We reason that the item to report is selected based on the precision with which its reported feature value is represented, and a swap error occurs when the location of that item is chosen incorrectly. We consider this to be more plausible than the possibility that a location is selected first based on the precision of the associated feature value, and then a different (lower precision) feature value from a different item is reported.

#### 2.2 Stochastic sampling model

The stochastic sampling model assumes that each memorized feature value is represented by a varying number of discrete samples with fixed precision. It can be derived as an idealization of neural population coding. The free parameters of this model are the sample precision $\omega_1$ and the mean total number of samples $\gamma$. The number of samples that contributes to the representation of each individual item is drawn independently from a Poisson distribution with mean $\gamma/N$. The resulting response distribution is then a mixture of von Mises distributions with different precisions, each corresponding to a certain number of samples and weighted with the probability of obtaining that sample count,

$$p(\hat{\theta}|\theta) = \sum_{k=0}^{\infty} \mathrm{Pr}_{\mathrm{Poisson}}\left(k; \frac{\gamma}{N}\right) \phi_\circ(\hat{\theta}; \theta, \kappa(k\omega_1)). \tag{2}$$

Here, $\mathrm{Pr}_{\mathrm{Poisson}}$ is the Poisson distribution,

$$\mathrm{Pr}_{\mathrm{Poisson}}(k; \lambda) = \frac{\lambda^k e^{-\lambda}}{k!}, \tag{3}$$

and $\phi_\circ$ is the von Mises distribution,

$$\phi_\circ(\hat{\theta}; \theta, \kappa) = \frac{1}{2\pi I_0(\kappa)} e^{\kappa\cos(\hat{\theta}-\theta)}. \tag{4}$$

The term $\kappa(\omega)$ is the concentration parameter that yields a von Mises distribution with precision $\omega$. With precision expressed as Fisher Information, the corresponding value

$\kappa$ can be obtained by numerically inverting the relationship $\omega = \kappa \frac{I_1(\kappa)}{I_0(\kappa)}$. $I_n$ is the modified Bessel function of the first kind. For fitting the model to data, we only consider sample counts $k$ within a range that covers the cumulative probabilities of the Poisson distribution from $10^{-5}$ to $1 - 10^{-5}$.

We note that the linear scaling of precision with the number of samples that we have assumed here is only an approximation of the exact distribution of ML estimates in circular space. We implemented the exact method [3] as a variant for all sampling models, and describe results of a comparison between the different variants in Section Full maximum likelihood decoding for circular feature spaces.

In the whole-report task with random response order, the stochastic sampling model predicts that there are no response correlations, because the number of samples is drawn independently for each item. The response distribution for this case is given by

$$p(\hat{\boldsymbol{\theta}}|\boldsymbol{\theta}) = \prod_{i=1}^{N} p(\hat{\theta}_i|\boldsymbol{\theta}), \tag{5}$$

where $p(\hat{\theta}_i|\boldsymbol{\theta})$ is the probability distribution including swap errors as defined in Eq. 1.

In the free response order condition, responses are ordered by precision, which is directly proportional to the number of samples that represent each item. This sorting induces positive correlations between response errors for consecutive responses within a trial. To compute the response probability for this condition, we determine all possible ordered sequences of sample counts $\boldsymbol{\lambda} = (\lambda_1, \ldots, \lambda_N), \lambda_i \geq \lambda_j \, \forall \, i < j$. The probability that each such sequence of sample counts will be generated by the model is

$$\Pr(\boldsymbol{\lambda}) = \prod_{k=0}^{\max(\boldsymbol{\lambda})} \Pr_{\text{Poisson}}\left(k; \frac{\gamma}{N}\right)^{n_k(\boldsymbol{\lambda})} \binom{N - \sum_{j=0}^{k-1} n_j(\boldsymbol{\lambda})}{n_k(\boldsymbol{\lambda})}, \tag{6}$$

where $n_k(\boldsymbol{\lambda})$ is the number of entries in $\boldsymbol{\lambda}$ with $\lambda_i = k$. The probability distribution for a sequence of responses (taking into account swap errors) is then

$$p(\hat{\boldsymbol{\theta}}|\boldsymbol{\theta}) = \sum_{\boldsymbol{\lambda}} \Pr(\boldsymbol{\lambda}) \prod_{i=1}^{N} \left( p_{\mathrm{T}} \phi_{\circ}(\hat{\theta}_i; \theta_i, \kappa(\lambda_i \omega_1)) + p_{\mathrm{NT}} \sum_{j \in \{1,\ldots,N\}, j \neq i} \phi_{\circ}(\hat{\theta}_i; \theta_j, \kappa(\lambda_i \omega_1)) \right). \tag{7}$$

For the implementation, we consider only sample counts $k$ with $\Pr_{\text{Poisson}}(k; \frac{\gamma}{N}) \geq 10^{-5}$, and we exclude the least likely sequences $\boldsymbol{\lambda}$ up to a cumulative probability of $10^{-3}$.

#### 2.3 Fixed sampling model

The fixed sampling model assumes that a fixed number $K$ of samples, each with a fixed precision $\omega_1$, is distributed as evenly as possible among the memory items in each trial. This model is mathematically equivalent to the slots+averaging model [4]. We note that the slot concept is commonly associated with an object- rather than feature-based view of working memory storage, but the sampling interpretation is agnostic with respect to this distinction.

The response probability distribution for the fixed sampling model is given by

$$p(\hat{\theta}|\theta) = \frac{K \bmod N}{N} \phi_{\circ}\left(\hat{\theta}; \theta, \kappa\left(\left\lceil \frac{K}{N} \right\rceil \omega_1\right)\right) + \left(1 - \frac{K \bmod N}{N}\right) \phi_{\circ}\left(\hat{\theta}; \theta, \kappa\left(\left\lfloor \frac{K}{N} \right\rfloor \omega_1\right)\right). \tag{8}$$

For the whole-report task, we again assume that responses are ordered by sample count. In the free response order condition, the response probability distribution is described by

$$p(\hat{\boldsymbol{\theta}}|\boldsymbol{\theta}) = \prod_{i=1}^{N} p(\hat{\theta}_i|\boldsymbol{\theta}) \tag{9}$$

with

$$p(\hat{\theta}_i|\theta) = \begin{cases} \phi_\circ\Big(\hat{\theta}_i; \theta, \kappa\big(\lceil \tfrac{K}{N} \rceil \omega_1\big)\Big), & \text{if } i \leq K \bmod N \\ \phi_\circ\Big(\hat{\theta}_i; \theta, \kappa\big(\lfloor \tfrac{K}{N} \rfloor \omega_1\big)\Big), & \text{otherwise.} \end{cases} \tag{10}$$

In the random response order condition, the fixed sampling model predicts negative correlations between response errors in a single trial (because having more samples for one item means fewer samples are available for others). We determine all possible sequences of sample counts $\boldsymbol{\lambda} = (\lambda_1, \ldots, \lambda_N), (\lambda_i = \lfloor \frac{K}{N} \rfloor \vee \lambda_i = \lceil \frac{K}{N} \rceil) \forall i, \sum_{i=1}^{N} \lambda_i = K$. Each such sequence will occur with equal probability $\Pr(\boldsymbol{\lambda}) = \left[\binom{N}{K \bmod N}\right]^{-1}$. The response probability $p(\hat{\boldsymbol{\theta}}|\boldsymbol{\theta})$ can then be expressed in the same way as in Eq. 7.

We tested the fixed sampling model (as well as the variant with random allocation described below) for values of $K$ in the range $(0, 25)$, going substantially beyond the typical estimates of three or four memory slots to ensure that model comparison results were not biased by a limited parameter range.

#### 2.4 Fixed sampling model with random allocation

As a variant of the fixed sampling model described above, we considered a model in which the total number of samples, $K$, is fixed, but each sample is randomly and independently assigned to one of the $N$ memory items with equal probability. The probability of obtaining a certain number $k$ of samples for a single item is then given by the binomial distribution,

$$\Pr_{\text{Bin}}\left(k; K, \frac{1}{N}\right) = \binom{K}{k} \left(\frac{1}{N}\right)^{k} \left(1 - \frac{1}{N}\right)^{K-k}, \tag{11}$$

and the response probability distribution is

$$p(\hat{\theta}|\theta) = \sum_{k=0}^{K} \Pr_{\text{Bin}}\left(k; K, \frac{1}{N}\right) \phi_\circ(\hat{\theta}; \theta, \kappa(k\omega_1)). \tag{12}$$

This model predicts correlations between response errors within a trial of the whole-report task in both the free response order and the random response order conditions. For the free response order condition, the possible sequences of ordered sample counts are $\{\boldsymbol{\lambda} | \sum_{i=1}^{N} \lambda_i = K, \lambda_i \geq \lambda_j \forall i < j\}$. The probability of each sequence can be determined as

$$\Pr(\boldsymbol{\lambda}) = \frac{1}{N^K} \prod_{i=1}^{N} \binom{K - \sum_{j=1}^{i-1} \lambda_j}{\lambda_i} \prod_{k=0}^{\max(\boldsymbol{\lambda})} \binom{N - \sum_{j=0}^{k-1} n_j(\boldsymbol{\lambda})}{n_k(\boldsymbol{\lambda})}. \tag{13}$$

For the random response order condition, the set of possible sample count sequences is $\{\boldsymbol{\lambda} | \sum_{i=1}^{N} \lambda_i = K\}$. The probability of each sequence is given by

$$\Pr(\boldsymbol{\lambda}) = \frac{1}{N^K} \prod_{i=1}^{N} \binom{K - \sum_{j=1}^{i-1} \lambda_j}{\lambda_i}. \tag{14}$$

The response probability $p(\hat{\boldsymbol{\theta}}|\boldsymbol{\theta})$ in the whole-report task can again be expressed as in Eq. 7.

#### 2.5 Stochastic sampling model with even allocation

A second model variant assumes that the total number of samples varies from trial to trial (as in the stochastic sampling model), but these samples are distributed across memory items as evenly as possible (as in the fixed sampling model). For each trial, the total number of samples is drawn from a Poisson distribution with mean $\gamma$. The probability distribution for a single response can then be given as weighted sum of probabilities from the fixed sampling model with different numbers of samples $k$,

$$p(\hat{\theta}|\theta) = \sum_{k=0}^{\infty} \mathrm{Pr}_{\mathrm{Poisson}}(k;\gamma) p_{\mathrm{fs}}(\hat{\theta}|\theta;k). \tag{15}$$

The response probabilities for the whole-report task can be determined in the same fashion as mixtures of the corresponding response probabilities in the fixed sampling model. We note that this introduces error correlations even in the case of the free response order condition, in which there are no correlations in the fixed sampling model.

#### 2.6 Generalized stochastic sampling model

In the generalized stochastic sampling model, the Poisson distribution over precision values is replaced by a negative binomial distribution with an additional discretization parameter $p$. The distribution of response errors is then given by

$$p(\hat{\theta}|\theta) = \sum_{k=0}^{\infty} \mathrm{Pr}_{\mathrm{NegBin}}\left(k; \frac{\gamma}{(1-p)N}, p\right) \phi_{\circ}(\hat{\theta};\theta,\kappa(k\omega_1 p)) \tag{16}$$

with

$$\mathrm{Pr}_{\mathrm{NegBin}}(k;r,p) = \frac{\Gamma(k+r)}{k!\,\Gamma(r)} p^r (1-p)^k \tag{17}$$

for $0 < p < 1$. For fitting the model to data, we only compute the sum over the most likely sample counts $k$ up to a cumulative probability of $1 - 10^{-4}$. We did not attempt to fit this model to whole-report data, as the number of combinatorial possibilities quickly becomes computationally infeasible as $p$ gets small.

#### 2.7 Gamma model

The Gamma model assumes that recall precision for each item is drawn independently from a Gamma distribution with shape parameter $\frac{\gamma}{N}$ and scale parameter $\omega_1$. This model constitutes the limit case of the generalized stochastic sampling model for $p \to 0$ (see Section Gamma distribution as limiting case), and has independently been proposed in two previous studies [5, 6]. In the formulation of van den Berg et al., the precision follows a Gamma distribution with mean $\bar{J}_1/N^{\alpha}$ and scale parameter $\tau$, which is identical to the model described here for $\bar{J}_1 = \gamma\omega_1$, $\tau = \omega_1$, and $\alpha = 1$ (see Section Power law for set size effects for model comparisons with $\alpha$ as free parameter).

The response probability distribution in the Gamma model is described as a continuous mixture of von Mises distributions,

$$p(\hat{\theta}|\theta) = \int_{\omega=0}^{\infty} p_{\text{Gamma}}\left(\omega; \frac{\gamma}{N}, \omega_1\right) \phi_\circ(\hat{\theta}; \theta, \kappa(\omega)) d\omega, \tag{18}$$

with

$$p_{\text{Gamma}}(\omega, k, \theta) = \frac{1}{\Gamma(k)\theta^k} \omega^{k-1} e^{-\frac{\omega}{\theta}}, \tag{19}$$

where $\Gamma$ is the gamma function. For model fitting, the integral is computed numerically with 1000 possible values of $\omega$, which cover the range of precision values with cumulative probabilities of the gamma distribution from $10^{-5}$ to $1 - 10^{-5}$.

In the whole-report task, the variable precision model predicts similar correlation patterns as the stochastic sampling model (which likewise draws precision values independently for each item). In the random response order condition, response errors within a trial are uncorrelated, and the precision distribution is given by

$$p(\hat{\boldsymbol{\theta}}|\boldsymbol{\theta}) = \prod_{i=1}^{N} p(\hat{\theta}_i|\boldsymbol{\theta}). \tag{20}$$

In the free response order condition, the probability distribution for a sequence of responses can be described as

$$p(\hat{\boldsymbol{\theta}}|\boldsymbol{\theta}) = \int_{\omega_1=0}^{\infty} \cdots \int_{\omega_N=0}^{\infty} p(\boldsymbol{\omega}) \prod_{i=1}^{N} \left( p_{\text{T}} \phi_\circ(\hat{\theta}_i; \theta_i, \kappa(\omega_i)) + p_{\text{NT}} \sum_{j \in \{1,\ldots,N\}, j \neq i} \phi_\circ(\hat{\theta}_i; \theta_j, \kappa(\omega_i)) \right) d\omega_1 \ldots d\omega_N, \tag{21}$$

where $\boldsymbol{\omega} = (\omega_1, \ldots, \omega_N)$ is the sequence of ordered precision values for the responses, and $p(\boldsymbol{\omega})$ is the probability of obtaining such a sequence if each precision value is drawn independently from a gamma distribution. Evaluating this equation is challenging, and in order to obtain an approximation, we discretize the range of possible precision values for each individual response into $m = 12$ bins of equal probability. We can then determine the probability of obtaining a sequence of ordered precision bin indices $\boldsymbol{b} = (b_1, \ldots, b_N)$ as

$$\Pr(\boldsymbol{b}) = \frac{1}{m^N} \prod_{k=1}^{m} \binom{N - \sum_{j=0}^{k-1} n_j(\boldsymbol{b})}{n_k(\boldsymbol{b})}. \tag{22}$$

Here, $n_k(\boldsymbol{b})$ denotes the number of entries in $\boldsymbol{b}$ with $b_i = k$, analogously to its use in Eq. 6. The response probability for a sequence of responses can then be given as weighted sum over all possible sequences $\boldsymbol{b}$,

$$p(\hat{\boldsymbol{\theta}}|\boldsymbol{\theta}) \approx \sum_{\boldsymbol{b}} \Pr(\boldsymbol{b}) \prod_{i=1}^{N} \int_{\omega=\omega_{\text{low}}(b_i)}^{\omega_{\text{high}}(b_i)} p_{\text{Gamma}}\left(\omega; \frac{\gamma}{N}, \omega_1\right) \left( p_{\text{T}} \phi_\circ(\hat{\theta}_i; \theta_i, \kappa(\omega)) + p_{\text{NT}} \sum_{j \in \{1,\ldots,N\}, j \neq i} \phi_\circ(\hat{\theta}_i; \theta_j, \kappa(\omega) \right. \tag{23}$$

Here, $\omega_{\text{low}}(b)$ and $\omega_{\text{high}}(b)$ are the boundaries of the precision bin with index $b$. Evaluating this form is more feasible, since the integrals over each precision bin can be computed independently (using the same numerical method with a total of 1000 sampling points as above) and then combined. We note that with this binning approach, we still obtain precise estimates of the response error distribution at each ordinal position; only the correlations between them are affected by the approximation.

#### 2.8 Neural population model with heterogeneous tuning curves

We tested a variant of the neural population model that incorporates heterogeneity in the cells' tuning functions of the kind observed in electrophysiological recordings (for the origin of this model and a more complete investigation of the topic, see [7]). Specifically, the model takes into account that neurons differ in their minimum (baseline) and maximum (peak) levels of activity, as well as in tuning width. The model also relaxes the assumption that the feature space is covered homogeneously by neural tuning curves. Instead, neurons' preferred values are selected at random from a uniform distribution. As in the original implementation of the neural population model [8], we assume that each of $N$ feature values in the memory sample array is encoded by a different population of $M$ neurons. The tuning curve of neuron $i$ encoding a feature value $\theta$ is given by a scaled von Mises distribution function plus a baseline,

$$f_i(\theta) = \alpha_i + \beta_i \exp\left(\kappa_i(\cos\left(\theta - \varphi_i\right) - 1)\right). \tag{24}$$

Here, $\alpha_i$ is the amplitude of the neuron's baseline activity, $\beta_i$ is the gain of the neuron, $\kappa_i$ is the von Mises concentration parameter which determines the tuning width, and $\varphi_i$ is the neuron's preferred feature value. These parameters are chosen randomly for each simulated neuron, with the degree of interneuron variability determined by a global heterogeneity parameter $\nu$.

For $\nu = 0$, the tuning parameters of all neurons are identical (with no baseline activity and homogeneous coverage of the feature space), making the model identical to the standard neural population model described in [8], and an exact circular analogue of the population model described in the main manuscript. The distributions of parameter values were chosen such that for $\nu = 1$, the population has approximately the heterogeneity observed in orientation-selective neurons in cortical area V1 [9]. For $\nu > 1$, individual neurons' parameters vary over wider ranges than observed in these biological populations.

Concretely, the parameters for each neuron are drawn from the following distributions:

$$\log\kappa_i \sim \mathcal{N}(\log\tilde{\kappa}, \nu^2) \tag{25}$$

$$\log\beta_i \sim \mathcal{N}(\log 1, \nu^2) \tag{26}$$

$$\log\alpha_i \sim \mathcal{N}(\log\left(0.04\nu\beta_i\right), 2) \tag{27}$$

$$\varphi_i \sim \mathcal{N}\left(\frac{2\pi}{M}(i-1), \nu^2\right) \bmod 2\pi \tag{28}$$

As in previous versions of the population model, we scaled the total expected activity of all neurons encoding all items with a population gain parameter, $\gamma$, which was fixed across changes in set size. However, in the heterogeneous model the information capacity of a neural population varied not only as a function of $\gamma$ but also all the individual tuning parameters of all the component neurons. In order to equate populations with different randomly-drawn tuning parameters, instead of treating $\gamma$ as a free parameter for model fitting, we instead used the expected precision of a decoded estimate as the free parameter, and set the population gain $\gamma$ to a value that would achieve it.

Specifically, we first normalized the tuning curves such that the population would on average fire exactly one spike within the decoding time interval:

$$\check{f}_i(\theta) = \frac{f_i(\theta)}{\frac{1}{2\pi}\sum_{j=1}^{M}\int_{-\pi}^{\pi} f_j(\theta)d\theta} \tag{29}$$

The mean precision of ML decoding from this population (assuming full knowledge of the tuning curves) was determined by the expected Fisher Information,

$$\check{\mathcal{I}} = \frac{1}{2\pi}\sum_{i=1}^{M}\int_{-\pi}^{\pi}\left(\frac{d\log \check{f}_i(\theta)}{d\theta}\right)^2 \check{f}_i(\theta)d\theta. \tag{30}$$

The mean decoding precision scales linearly with the number of spikes available for decoding, so in order to achieve the desired precision $\bar{\mathcal{I}}$, we set the global gain parameter $\gamma$ to

$$\gamma = \bar{\mathcal{I}}/\check{\mathcal{I}}. \tag{31}$$

The spike count $r_i$ of neuron $i$ encoding feature value $\theta$ in a trial with set size $N$ was then drawn from a Poisson distribution,

$$r_i \sim \mathrm{Poisson}\Big(\frac{\gamma}{N}\check{f}_i(\theta)\Big). \tag{32}$$

The log likelihood of stimulus feature $\theta'$ for a given set of spikes $\boldsymbol{r}$ is (up to addition by a constant),

$$\log \mathcal{L}(\theta'|\boldsymbol{r}) = \sum_{i=1}^{M} r_i \log \check{f}_i(\theta') - \check{f}_i(\theta'). \tag{33}$$

Decoded estimates were obtained as the maximum of this function, and their precision as the width of the likelihood function measured in terms of Fisher Information.

The heterogeneous model therefore has three free (global) parameters that determine the distributions of the single neuron parameters and thereby the predicted error distributions of decoded estimates: the median tuning curve width, $\tilde{\kappa}$, the heterogeneity parameter, $\nu$, and the mean precision for a single stored item, $\bar{\mathcal{I}}$. We estimated the response distributions for this model by sampling. For each combination of values for the parameters $\tilde{\kappa}$, $\nu$, and $\bar{\mathcal{I}}$ on a search grid (described in Section Fitting procedure), we randomly drew 100 sets of single-neuron parameters for $M$ = 1000 neurons[1] from the distributions specified in Eqs. 25 – 28. For each set of single-neuron parameters, we generated a neural spiking pattern in response to 1000 randomly chosen feature values $\theta$, and obtained likelihood functions and ML estimates $\hat{\theta}$ as described above. The response error distribution is then approximated by a histogram over the decoding errors, $\hat{\theta} \ominus \theta$, averaged over all sets of single-neuron parameters ($\ominus$ indicates subtraction on the circle). Mean ML parameters obtained for the single-report dataset were $\tilde{\kappa}$ = 1.53 $\pm$ 0.15, $\bar{\mathcal{I}}$ = 18.6 $\pm$ 1.0, and $\nu$ = 0.66 $\pm$ 0.08.

Unlike the discrete distribution predicted by the (homogeneous) stochastic sampling model (Fig. S6E), there is no probability of zero precision decoding. This is because the unevenness in coverage of the stimulus space makes no spikes a more probable response to some feature values than others, meaning it is no longer uninformative about the stimulus. However, at lower set sizes there is a sharp increase in probability of very low precision estimates that could not in practice be discriminated from zero (blue curve in Fig. S6D).

Example precision distributions from the Gamma (variable precision; [5, 6]) model are shown in Fig. S6F. Interestingly, heterogeneity provides a second putative connection between population coding and Gamma-distributed precision, in addition to the

[1]The precise number of neurons has very little influence on the response distributions once the average spacing between neurons' preferred values is significantly smaller than the tuning curve width, and therefore we do not treat $M$ as a free parameter in this model.

one set out in the main text. This is because a Gamma process (the random process whose marginal distribution at each moment in time is a Gamma distribution) can be constructed from an infinite superposition of different Poisson processes, varying in their rate and, inversely, in their amplitude (i.e. Lévy jump size). So, with the right kind of heterogeneity, a population model with Poisson spiking could theoretically result in estimates with exactly Gamma-distributed precision.

We also fit a further variant of the heterogeneous model incorporating short-range correlations of the form

$$c_{ij} = c_0 \exp(- |\varphi_i \ominus \varphi_j|), \tag{34}$$

with $c_0$ set to 0.2. We assumed that the decoder did not have knowledge of the correlation structure. Due to the significant computational challenge of simulating correlated Poisson activity, we used a Gaussian approximation to Poisson (see [7] for details), and compared model fits to an otherwise identical model without correlations (i.e. with $c_0$ set to 0).

### 3 Fitting procedure

We fit models separately to the behavioral data of each participant in each experiment of the single-report and whole-report dataset. Data of each participant across all set sizes was fit with a single set of parameter values. We employed two different methods to determine the ML parameter values, namely the Nelder-Mead simplex algorithm and grid search over the parameter space.

We used the Nelder-Mead simplex algorithm to fit all models except for the generalized stochastic model and the neural population model with heterogeneous tuning curves, for both single-report and whole-report data. We defined a limited grid of initial parameter values, and ran the fitting algorithm (function *fminsearch* in Matlab) with each possible combination of initial values until a termination tolerance of 0.01 was reached for both the fitted parameter values and the resulting likelihood value. Possible initial values for the sample precision $\omega_1$ were $2^0, 2^2, 2^4$. For stochastic sampling models and gamma model, we first defined initial values for the mean precision at set size one, $E[\omega]$, as $2^2, 2^4, 2^6$, then determined initial values of $\gamma$ as $\gamma = \frac{E[\omega]}{\omega_1}$. For fixed sampling models, we obtained separate fits for all integer values of $K$ in the range $(0, 25)$, and selected the fit with the highest likelihood. Initial values for $p_{\mathrm{NT}}$ were $0.01, 0.05, 0.1$, and for $\alpha$ they were $2^{-0.5}, 2^0, 2^{0.5}$. In variants where these parameters were not used they were fixed at $p_{\mathrm{NT}} = 0$ and $\alpha = 1$, respectively.

We used the grid search to fit the generalized stochastic model (separately for different values of the discretization parameter $p$) and the neural population model with heterogeneous tuning curves to single-report data. For the latter, likelihood values were determined by Monte Carlo sampling, as no closed form solution is available. We also obtained additional fits for the models described in the main text (stochastic sampling, fixed sampling, random-fixed, even-stochastic, and gamma) to verify that the Nelder-Mead simplex algorithm for these models terminated in the global rather than a local maximum of the likelihood function. Results shown in Fig. 3E of the main manuscript are based on the grid search fits to allow fair comparison between the stochastic sampling (Poisson) model, generalized stochastic sampling model, and Gamma model.

The parameter grid was spanned by 50 possible values of each model parameter. Values for sample precision, $\omega_1$, were spaced logarithmically in the range $[2^{-4}, 2^5]$, and values for mean precision, $E[\omega]$, in the range $[2^{-2}, 2^9]$. For the fixed sampling models, the parameter $K$ took all integer values in the range $[0, 50)$. The values for proportion of non-target responses, $p_{\mathrm{NT}}$, were evenly spaced in the range $[0.0, 0.14]$ for all models. The values for the discretization parameter, $p$, in the generalized stochastic sampling model was spaced logarithmically in the range $[10^{-4}, 1]$. To compute likelihood values in the grid search, response errors were discretized into 101 evenly spaced bins for all models as well as for the behavioral data (ensuring fair comparison between models with closed-form likelihood function and the heterogeneous neural model which requires sampling).

In both fitting methods, we determined a maximum likelihood value $L$ and an associated set of parameters. For comparison between models that differed in the number of free parameters, we computed Akaike information criterion (AIC) scores,

$$\mathrm{AIC} = 2k - 2\log(L), \tag{35}$$

and Bayesian information criterion (BIC) scores,

$$\mathrm{BIC} = \log(n)k - 2\log(L). \tag{36}$$

Here, $k$ is the number of free parameters in each model, and $n$ is the number of data points (number of trials in the single-report tasks, and number of individual responses in the whole-report tasks). AIC and BIC differences for the models described in the main text are depicted in Fig. S3A and E for single-report and whole-report data, respectively. ML fit values of free parameters are reported in Tables S3 and S4.

### 4 Additional model comparisons

#### 4.1 Full maximum likelihood decoding for circular feature spaces

The assumption that the decoding precision in sampling models increases in equal discrete steps with the number of samples is only strictly true for certain cases. For circular feature spaces with samples drawn from a von Mises distributions, it is only an approximation. An exact method to compute the distribution of response errors arising from ML decoding in circular space was derived in [8] and [3]. For a given number of samples, $m$, that are drawn independently from the same von Mises distribution with concentration parameter $\kappa_1 = \kappa(\omega_1)$, the resulting distribution of decoding error can be described as a continuous scale mixture of von Mises distributions,

$$p(\hat{\theta}|\theta, m) = \int p(r|m, \kappa_1)\phi_\circ(\hat{\theta}; \theta, r\kappa_1)dr, \tag{37}$$

with

$$p(r|m, \kappa) = \frac{I_0(\kappa r)}{(I_0(\kappa))^m} r\psi_m(r). \tag{38}$$

Here, $r\psi_m(r)$ is the probability density function for the resultant length $r$ of a uniform random walk of $m$ steps. The distribution of response errors in each sampling model is then a mixture of probability distributions $p(\hat{\theta}|\theta, m)$, weighted with the probability of obtaining $m$ samples for an item.

We obtained ML fits using this method to determine response error distributions for the stochastic sampling model, fixed sampling model, and random-fixed and even-stochastic variants (the method is not compatible with the Gamma model, since this model does not use discrete samples). The quality of fit was improved for all models (Fig. S3B and F), with only minimal changes for the stochastic sampling model, and largest improvements for fixed sampling model and random-fixed model fits to single-report data. However, the overall pattern of results did not change when employing exact ML decoding instead of the simpler approximation.

Mean parameter values for the models with exact ML decoding differ substantially from those obtained using the approximation method. This is driven by a subset of participants for which the best model fits using the exact method are characterized by a large number of low-precision samples. Within this region of the parameter space, the exact method and the approximation deviate more strongly from each other (Fig. S7), with the exact method producing slightly broader distributions of response errors that can provide a better fit to some experimental result. We note that for the fixed sampling model, the variant with exact ML decoding produces fits with $K > 10$ for a substantial proportion of participants (40% in the single-report dataset, 18% in the whole-report dataset), which is in conflict with typical estimates of 3-4 memory slots.

#### 4.2 Excluding swap errors

We obtained ML fits of the behavioral data for all models without swap errors by keeping the parameter $p_{\mathrm{NT}}$ fixed at zero. The quality of fit for all models decreased substantially in this variant, independent of whether we measured it via AIC or BIC values (which differ in how strongly they penalize additional free parameters; Fig. S3C and G). This is consistent with previous findings that inclusion of swap errors improves model fit [1, 10].

#### 4.3 Power law for set size effects

The variable precision model [5] proposed that the effect of set size on mean recall precision is best explained by a power law of the form $E[\omega] \propto N^{-\alpha}$, with a free parameter $\alpha$. We added this parameter to the formulations of the stochastic sampling model and the Gamma model (the other models assume that a certain number of samples is distributed between all items, thus the power law is not readily applicable). Quality of fit was improved for both models (Fig. S3D and E), with the stochastic sampling model still providing better quality of fit for both single-report and whole-report datasets.

#### 4.4 Random drift over memory delays

The most striking effect in the whole-report data [2] is the salient decrease in recall precision for successive responses when response order is freely chosen. A weaker decrease in precision, however, can also be observed in the random response order condition (Fig. S4B), where it cannot be explained by a strategy of reporting the best-remembered items first. All models described so far lack any mechanism that could capture the effects of increasing memory delay for later responses, or interference from intervening reports. Thus, they inevitably fail to fit this aspect of the behavioral data (the model fits for successive responses at each set size are identical in Fig. S4B).

We tested a mechanism for random drift of memorized feature values during memory delays that was previously described for the population coding model [11]. The effect of this drift is described by convolving the response probability distribution of each model with a wrapped normal distribution centered on zero,

$$\tilde{p}(\hat{\theta}) = (p * f_{\mathrm{WM}}(0, \sigma))(\hat{\theta}). \tag{39}$$

We assume that the parameter $\sigma$ increases linearly from the first to the last response in each trial of the whole-report task. Moreover, we assume that the rate of random drift also scales linearly with set size, which has been found to provide the best fits to delayed reproduction data with varying memory delays [11]. This yields

$$\sigma_i = iN\eta \tag{40}$$

for the $i$th response in a whole-report trial, where $\eta$ is a new free parameter specifying the base drift rate.

When fitting a model with random drift to whole-report data, there is the concern that a high drift rate provides an alternative mechanism to produce the salient decrease in precision observed in the free response order condition. The much weaker decrease in precision in the random response order should constrain the drift rate parameter, but the two conditions were performed by separate groups of participants. To see whether a single drift rate parameter can account for response patterns across response order conditions, we pooled participant data to create a single super-participant each for the color report (Experiments 1a & 2a) and orientation report conditions (Experiments 1b & 2b). We then fit this data with one set of parameters each, varying only whether responses were ordered by memory precision or not to capture the different response order conditions (Table S5).

Pooling the data did not qualitatively change the pattern of quality-of-fit measures for the whole-report task (Fig. S3I), and the inclusion of a drift rate improved quality of fits in all models (Fig. S3J). The stochastic sampling model still provides the best fit for the data, and it closely reproduces the overall pattern of response distributions across response order conditions (Fig. S5), including the nearly uniform response distributions for the last reports at set size 6. We note that the slots+averaging model may be at a disadvantage in fitting this pooled data since it is forced to commit to a fixed number of slots, whereas the data may come from participants with different numbers of slots. For a more fair and thorough model comparison, it would desirable to obtain data from individual participants performing both free and random response order conditions of the whole-report task.

### 5 Limiting cases of the negative binomial distribution

In the generalized stochastic sampling model, precision follows a scaled negative binomial distribution,

$$\frac{\omega}{\omega_1 p} \sim \mathrm{NegBin}\left(\frac{\gamma}{N(1-p)}, p\right), \tag{41}$$

with parameters $\gamma > 0$, $\omega_1 > 0$, and $0 < p < 1$. The probability of a precision value $\omega$ that is an integer multiple of $\omega_1 p$ is defined as

$$\Pr(\omega) = \frac{\Gamma(\frac{\omega}{\omega_1 p} + \frac{\gamma}{N(1-p)})}{(\frac{\omega}{\omega_1 p})!\,\Gamma(\frac{\gamma}{N(1-p)})} p^{\frac{\gamma}{N(1-p)}} (1-p)^{\frac{\omega}{\omega_1 p}} \quad \text{for} \quad \frac{\omega}{\omega_1 p} \in \mathbb{Z}^{\geq 0}. \tag{42}$$

All other precision values have zero probability.

This probability distribution has mean $E[\omega] = \gamma\omega_1/N$ and variance $Var[\omega] = \gamma{\omega_1}^2/N$. Interpreted as a discrete sampling model, the expected number of samples per item is $\gamma/(pN)$ with variance $\gamma/(p^2 N)$.

#### 5.1 Convergence result for large N

For a set size of $N$, the number of items with precision exceeding a threshold, defined as a multiple $m > 0$ of the base precision $\omega_1$, follows a binomial distribution,

$$S_{\omega > m\omega_1} \sim \mathrm{Bin}(N, \Pr(\omega > m\omega_1)), \tag{43}$$

with mean

$$E[S_{\omega > m\omega_1}] = N \Pr(\omega > m\omega_1). \tag{44}$$

For the generalized stochastic sampling model, with precision following a scaled negative binomial distribution, we have

$$\begin{aligned} N \Pr(\omega > m\omega_1) &= N \sum_{k=\lfloor m/p+1 \rfloor}^{\infty} \frac{\Gamma(k + \frac{\gamma}{N(1-p)})}{k!\,\Gamma(\frac{\gamma}{N(1-p)})} p^{\frac{\gamma}{N(1-p)}} (1-p)^k \\ &= N \frac{p^{\frac{\gamma}{N(1-p)}}}{\Gamma\left(\frac{\gamma}{N(1-p)}\right)} \sum_{k=\lfloor m/p+1 \rfloor}^{\infty} \frac{\Gamma(k + \frac{\gamma}{N(1-p)})}{k!} (1-p)^k. \end{aligned} \tag{45}$$

In the limit $N \to \infty$, we have

$$p^{\frac{\gamma}{N(1-p)}} \to 1, \tag{46}$$

$$\frac{\Gamma\left(k + \frac{\gamma}{N(1-p)}\right)}{k!} \to \frac{\Gamma(k)}{k!} = \frac{1}{k} \quad \text{for} \quad k \in \mathbb{Z}^{\geq 0}, \tag{47}$$

and

$$\frac{N}{\Gamma\left(\frac{\gamma}{N(1-p)}\right)} \to \frac{\gamma}{1-p}. \tag{48}$$

We obtain

$$\lim_{N \to \infty} E[S_{\omega > m\omega_1}] = \gamma \sum_{k=\lfloor m/p+1 \rfloor}^{\infty} \frac{(1-p)^{k-1}}{k}. \tag{49}$$

Note that all terms of the sum are strictly positive. Convergence of the sum can be demonstrated by a ratio test. The ratio of successive terms is

$$R = \frac{(1-p)^k k}{(1-p)^{k-1}(k+1)} = (1-p)\frac{k}{k+1}, \tag{50}$$

so

$$\lim_{k \to \infty} |R| = 1 - p < 1, \tag{51}$$

as $p > 0$. So the mean number of items with above-threshold precision converges to a finite positive number at large set sizes, for all model parameters, implementing a probabilistic form of “item limit”.

#### 5.2 Poisson distribution as limiting case

To determine the limiting cases of the negative binomial distribution for $p \to 1$ and $p \to 0$, we use its characteristic function. For $X \sim \mathrm{NegBin}(r, p)$, this is given by

$$\varphi_\omega(t) = E[e^{itX}] = \left(\frac{p}{1-(1-p)e^{it}}\right)^r. \tag{52}$$

For the scaled negative binomial distribution as used in the generalized stochastic model (Eq. 41), the characteristic function is

$$\varphi_\omega(t) = \left(\frac{p}{1-(1-p)e^{it\omega_1 p}}\right)^{\frac{\gamma}{N(1-p)}}. \tag{53}$$

We can write this as

$$\begin{aligned}\varphi_\omega(t) &= \exp\left[\log\left[\left(\frac{p}{1-(1-p)e^{it\omega_1 p}}\right)^{\frac{\gamma}{N(1-p)}}\right]\right] \\ &= \exp\left[\frac{\gamma}{N}\left(\frac{\log(p)}{1-p} - \frac{\log(1-(1-p)e^{it\omega_1 p})}{1-p}\right)\right]\end{aligned} \tag{54}$$

In the limit $p \to 1$, we have

$$\frac{\log(p)}{1-p} \to -1 \tag{55}$$

and for $q = 1-p$ (and thus $q \to 0$)

$$\frac{\log(1+aq)}{q} \to a. \tag{56}$$

The limiting case of the characteristic function for $p \to 1$ is therefore

$$\varphi_\omega(t) \to \exp\left[\frac{\gamma}{N}(e^{it\omega_1} - 1)\right], \tag{57}$$

which is the characteristic function of the scaled Poisson distribution, $\frac{\omega}{\omega_1} \sim \mathrm{Poisson}(\frac{\gamma}{N})$.

#### 5.3 Gamma distribution as limiting case

We again start with characteristic function of the scaled negative binomial distribution (Eq. 53). We use the Taylor series for the exponential function,

$$e^x = 1 + x + \mathcal{O}(x^2) \quad \text{as} \quad x \to 0, \tag{58}$$

where $\mathcal{O}(x^2)$ designates terms that are of order $x^2$ or greater.

We can then write the characteristic function as

$$\begin{aligned}\varphi_\omega(t) &= \left(\frac{p}{1-(1-p)(1+it\omega_1 p + \mathcal{O}(p^2))}\right)^{\frac{\gamma}{N(1-p)}} \\ &= \left(\frac{p}{p(1-it\omega_1) + \mathcal{O}(p^2)}\right)^{\frac{\gamma}{N(1-p)}} \\ &= \left(\frac{1}{1-it\omega_1 + \mathcal{O}(p)}\right)^{\frac{\gamma}{N(1-p)}}.\end{aligned} \tag{59}$$

In the limit $p \to 0$, this yields

$$\varphi_\omega(t) \to \left(\frac{1}{1 - it\omega_1}\right)^{\frac{\gamma}{N}}. \tag{60}$$

This is the characteristic function of the Gamma distribution with $\omega \sim \mathrm{Gamma}(\frac{\gamma}{N}, \omega_1)$. We are grateful to Zakhar Kabluchko for this derivation.

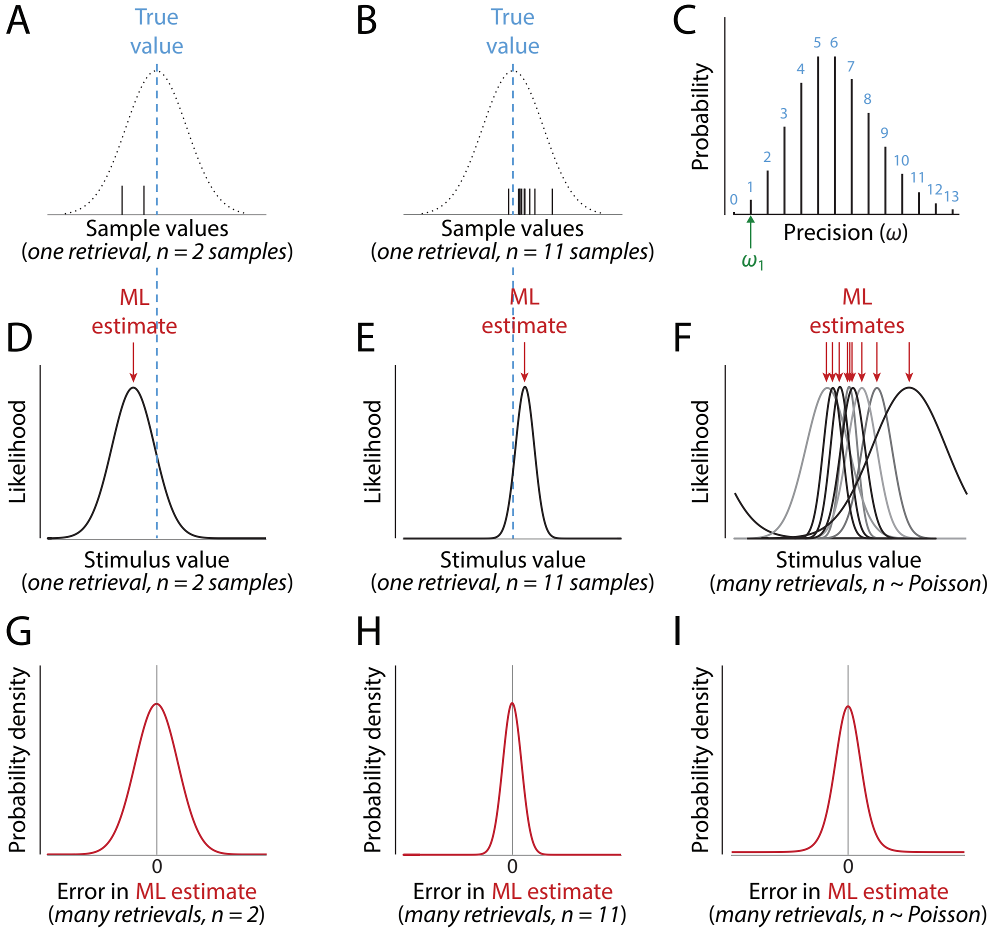


Figure S1: Likelihood and variability in the stochastic sampling model. (A, B) Generation of samples. Retrieval of a stimulus feature corresponds to maximum likelihood (ML) estimation based on a set of $n$ samples, each drawn from a normal distribution centered on the true feature value with a fixed precision (inverse of variability), $\omega_1$. The number of samples is drawn from a Poisson distribution. Examples shown are of estimation for $n$ = 2 in (A) and panels below; $n$ = 11 in (B) and panels below. (C) From retrieval to retrieval, precision (measured with respect to the likelihood) varies with a scaled Poisson distribution, taking on discrete values corresponding to the number of samples (blue) multiplied by the sample precision $\omega_1$. (D, E) The likelihood function on any particular retrieval measures the compatibility of different stimulus values with the obtained samples. For normally distributed samples, the likelihood is also normal with peak (the ML estimate) at the mean of the sample values. The likelihood width (corresponding to precision $n\omega_1$) indicates the reliability of the estimate and predicts subjective confidence. (F) Both the location and width of the likelihood vary between retrievals. (G, H) Considering only retrievals based on $n$ samples, the error in the ML estimate varies according to a normal distribution with precision $n\omega_1$, equal to the precision of the corresponding likelihood functions. (I) Over all retrievals, the estimation error has a non-normal (leptokurtic) distribution, because it comprises a mixture of normal distributions with different precisions (as in (C)).

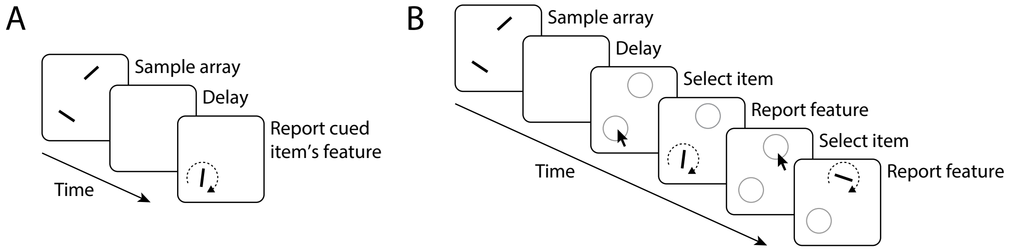


Figure S2: Delayed reproduction tasks. (A) Structure of an illustrative single-report task with orientation report cued by object location. (B) Structure of an illustrative whole-report task. The participant sequentially reports the features of all items in the sample array, either in a freely chosen order (shown here) or a prescribed random order.

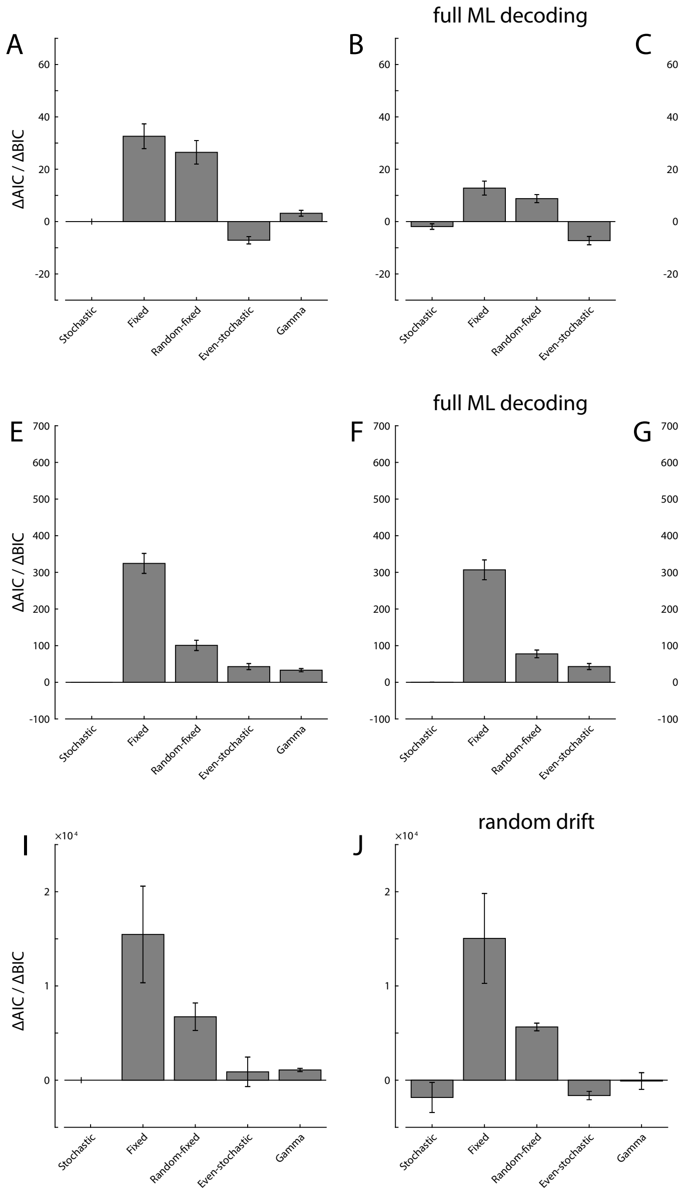


Figure S3: Model comparison for additional model variants. Mean differences in AIC (light gray) and BIC values (dark gray) relative to the stochastic sampling model with swap errors are shown for single-report data (A-D), whole-report data (E-H), and whole-report data pooled over participants (I-J). Better models have lower values. Error bars indicate $\pm$ 1 SE. (A, E, I) Models as described in the main text, including a fixed probability of swap errors per memory item. (B, F) Variant of discrete sampling models with exact ML decoding from samples in circular feature space. (C, G) Model variants without swap errors ($p_{\mathrm{NT}}$ = 0). (D, H) Model variants with power law relationship for set size effect, with exponent $\alpha$ as additional free parameter. (J) Model variants with random drift of memorized features over memory delay.

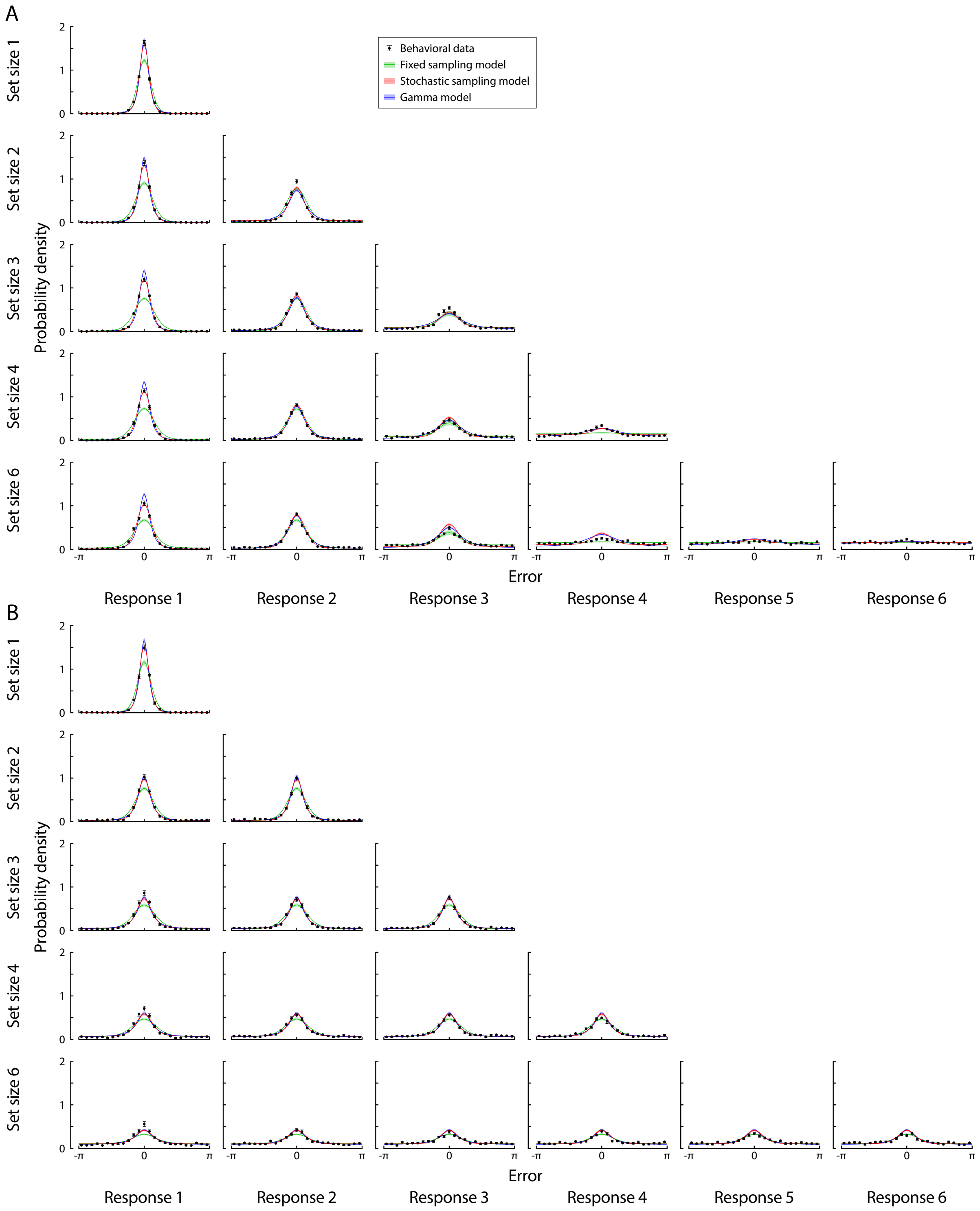


Figure S4: Behavioral data and model fits in the whole-report task [2] for color. (A) Free response order condition (Experiment 1a). (B) Random response order condition (Experiment 2a). Solid lines show the mean across participants, and error bars and shaded areas indicate $\pm$ 1 SE.

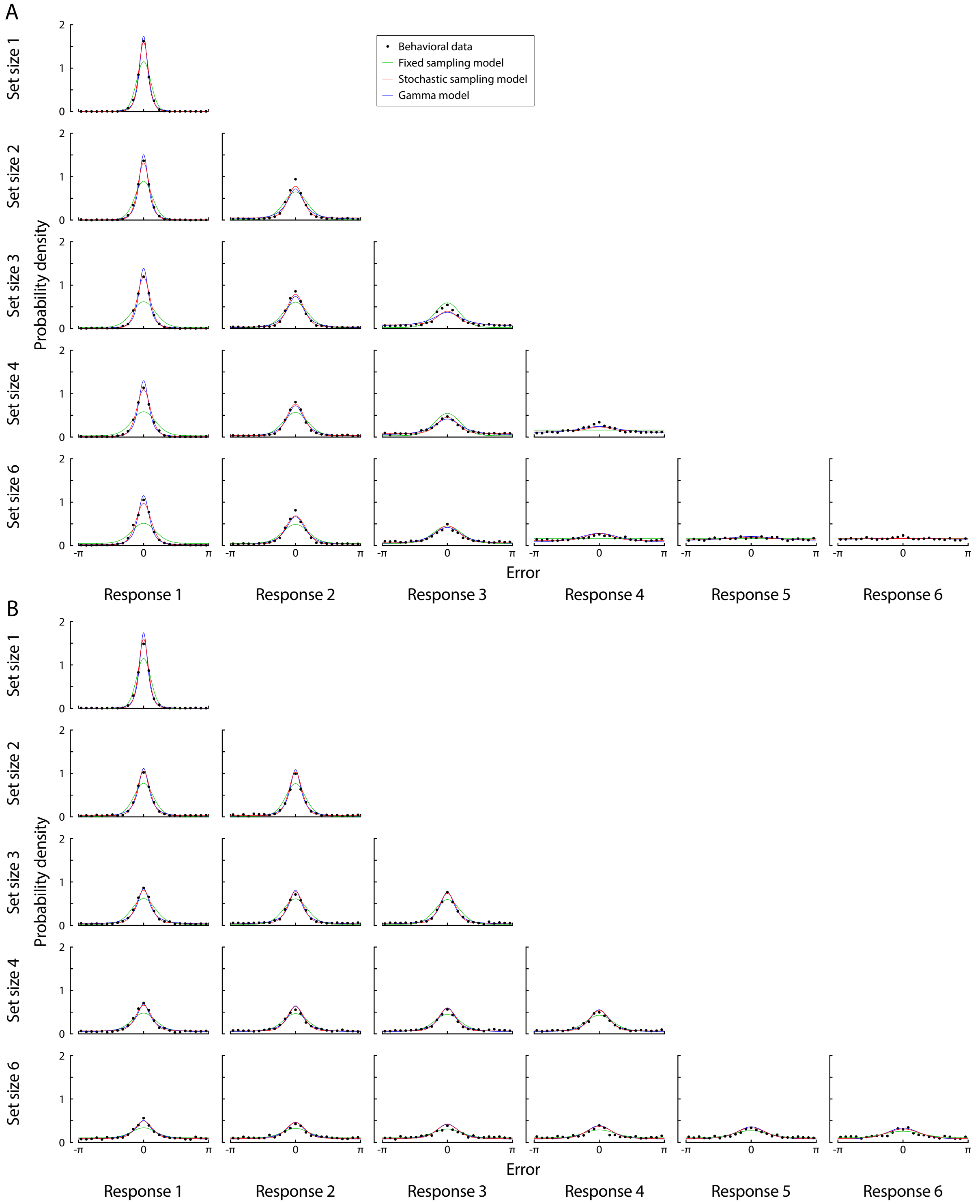


Figure S5: Behavioral data pooled across participants and model fits with random drift of memorized feature values. Panels as in Fig. S4. For each model, a single set of parameters was obtained by ML fit across the two response order conditions, varying only in whether responses are ordered by precision of memory representations.

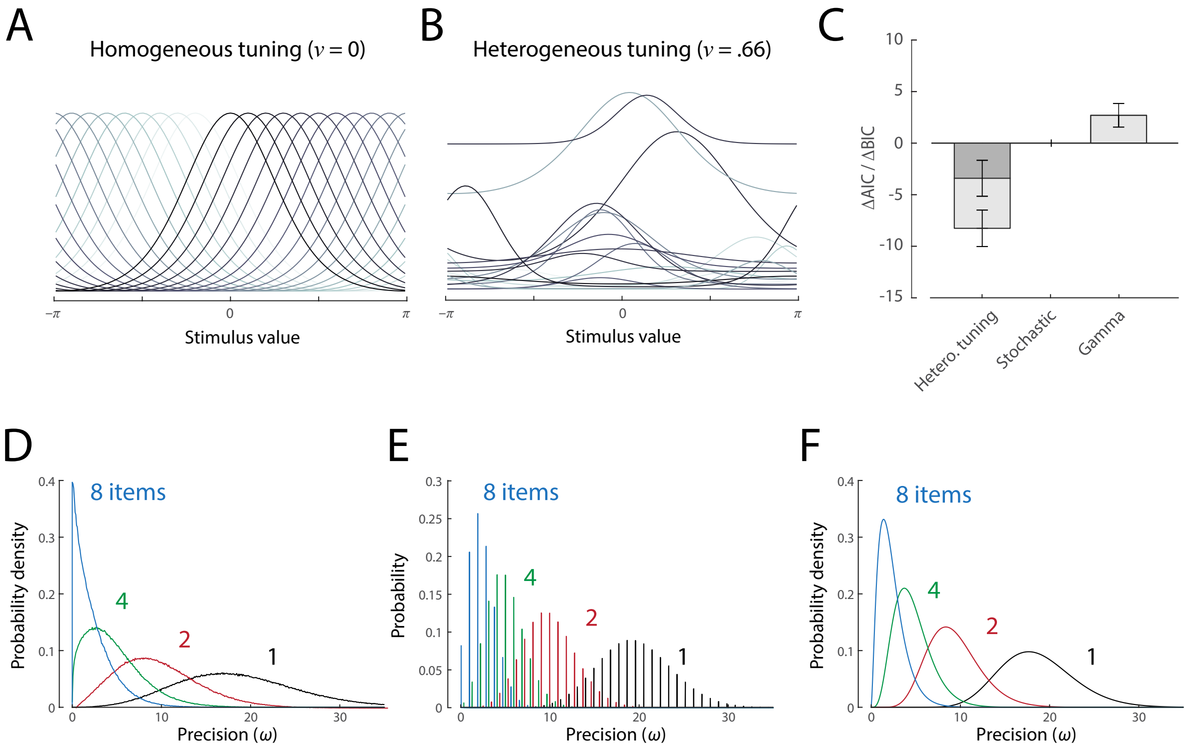


Figure S6: Population coding model with heterogeneous tuning. (A) Examples of homogeneous tuning functions in the idealized neural population underlying the stochastic sampling model. (B) Examples of heterogeneous tuning functions corresponding to mean parameters of the fitted heterogeneous model. (C) Results of model comparison on single-report data. The heterogeneous model out-performed the stochastic sampling model, and the model with Gamma-distributed precision, according to both AIC and BIC measures. (D) Distributions of precision of decoded estimates in the heterogeneous model, for different set sizes, based on mean parameters of best fit. (E) Distributions of precision for the stochastic sampling model with matched parameters, for comparison. (F) Distributions of precision for the Gamma model with matched parameters.

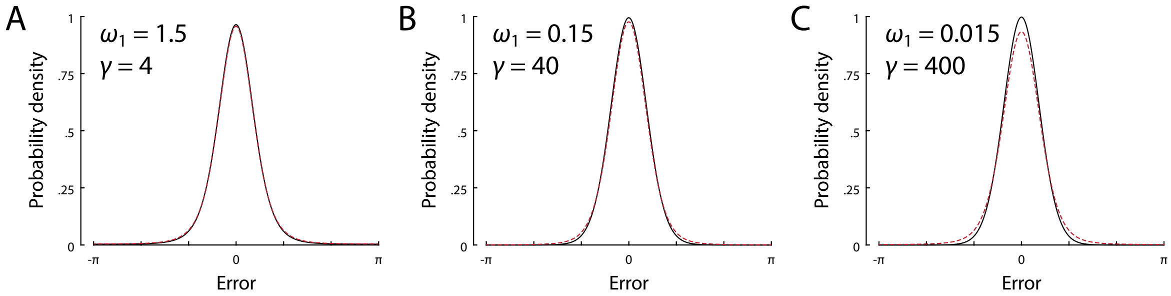


Figure S7: Distributions of response errors in the stochastic sampling model, using either exact ML decoding in circular space (red dashed lines) or the approximation assuming a linear increase in precision (expressed as Fisher information) with the number of samples (black lines). The approximation is very close for typical parameter values (A; parameters here are close to median values of the stochastic sampling model fits scaled for set size 4), but deviations increase when moving to parameterizations with a higher number of low-precision samples (B, C).

Table S1: Single-report experiments used for model comparison in this study. The Trials column denotes the number of trials each participant completed per set size.

| No | Study | Feature | Set Sizes | Participants | Trials |
|---|---|---|---|---|---|
| 1 | Zhang & Luck, 2008 [4] | Color | 1, 2, 3, 6 | 8 | 125 |
| 2 | Bays et al., 2009 [10] | Color | 1, 2, 4, 6 | 12 | 200 |
| 3 | van den Berg et al., 2012 [5] | Color | 1 – 8 | 13 | 216 |
| 4 | van den Berg et al., 2012 [5] | Orientation | 1 – 8 | 6 | 320 |
| 5 | Rademakers et al., 2012 [12] | Orientation | 3, 6 | 6 | 800 |
| 6 | Bays, 2014 [8], Exp 1 | Orientation | 1, 2, 4, 8 | 8 | 230 |
| 7 | Bays et al., 2011 [13], Exp 1 | Orientation | 1, 2, 4, 6 | 8 | 800 |
| 8 | Bays, Wu & Husain, 2011 [14] | Orientation | 1, 6 | 10 | 50, 250 |
| 9 | Bays, Wu & Husain, 2011 [14] | Color | 1, 6 | 10 | 50, 250 |
| 10 | Gorgoraptis et al., 2011 [15] | Orientation | 1 – 5 | 8 | 100 |
| 11 | Pratte et al., 2017 [16] | Orientation | 1, 2, 3, 6 | 12 | 640 |

Table S2: Whole-report experiments used for model comparison. All experiments are taken from [2].

| No | Feature | response order | Set Sizes | Participants | Trials |
|---|---|---|---|---|---|
| 1 | Color | Free | 1, 2, 3, 4, 6 | 22 | 99 |
| 2 | Orientation | Free | 1, 2, 3, 4, 6 | 20 | 200 |
| 3 | Color | Random | 1, 2, 3, 4, 6 | 17 | 99 |
| 4 | Orientation | Random | 1, 2, 3, 4, 6 | 19 | 200 |

Table S3: Parameter values of ML fits for single-report data (mean $\pm$ 1 SE across participants and experiments). Outliers with deviation from mean greater than 3 SD were excluded (at most 4 out of 101 individual fit values for each parameter) in order to provide more representative parameter values.

| Model | $\gamma$ or $K$ | $\omega_1$ | $p_{\mathrm{NT}}$ | $\alpha$ |
|---|---|---|---|---|
| Stochastic | 13.2 $\pm$ 1.7 | 1.84 $\pm$ 0.11 | 0.0245 $\pm$ 0.0023 | |
| Fixed | 4.90 $\pm$ 0.21 | 2.80 $\pm$ 0.14 | 0.0265 $\pm$ 0.0023 | |
| Random-fixed | 11.4 $\pm$ 0.7 | 1.55 $\pm$ 0.10 | 0.0238 $\pm$ 0.0023 | |
| Even-stochastic | 7.31 $\pm$ 1.47 | 3.04 $\pm$ 0.15 | 0.0287 $\pm$ 0.0024 | |
| Gamma | 8.63 $\pm$ 1.94 | 5.00 $\pm$ 0.36 | 0.0281 $\pm$ 0.0024 | |
| *full ML decoding for circular feature spaces* | | | | |
| Stochastic | 35.0 $\pm$ 6.3 | 1.68 $\pm$ 0.12 | 0.0261 $\pm$ 0.0024 | |
| Fixed | 11.7 $\pm$ 0.9 | 2.10 $\pm$ 0.17 | 0.0305 $\pm$ 0.0027 | |
| Random-fixed | 13.8 $\pm$ 0.8 | 1.46 $\pm$ 0.11 | 0.0262 $\pm$ 0.0024 | |
| Even-stochastic | 15.9 $\pm$ 3.3 | 2.93 $\pm$ 0.17 | 0.0289 $\pm$ 0.0024 | |
| *excluding swap errors* | | | | |
| Stochastic | 8.35 $\pm$ 0.81 | 2.28 $\pm$ 0.13 | | |
| Fixed | 3.67 $\pm$ 0.13 | 3.25 $\pm$ 0.17 | | |
| Random-fixed | 6.91 $\pm$ 0.34 | 2.08 $\pm$ 0.12 | | |
| Even-stochastic | 4.49 $\pm$ 0.14 | 3.55 $\pm$ 0.16 | | |
| Gamma | 3.95 $\pm$ 0.73 | 8.16 $\pm$ 0.62 | | |
| *power law for set size effects* | | | | |
| Stochastic | 8.46 $\pm$ 0.51 | 1.96 $\pm$ 0.11 | 0.0310 $\pm$ 0.0026 | 0.809 $\pm$ 0.031 |
| Gamma | 4.35 $\pm$ 0.37 | 5.60 $\pm$ 0.40 | 0.0339 $\pm$ 0.0026 | 0.809 $\pm$ 0.032 |

Table S4: Parameter values of ML fits for whole-report data (mean ± 1 SE across participants and experiments). Outliers with deviation from mean greater than 3 SD were excluded (at most 4 out of 78 individual fit values for each parameter).

| Model | $\gamma$ or $K$ | $\omega_1$ | $p_{\mathrm{NT}}$ | $\alpha$ |
|---|---|---|---|---|
| Stochastic | 4.30 ± 0.14 | 5.18 ± 0.25 | 0.0367 ± 0.0041 | |
| Fixed | 2.72 ± 0.10 | 4.85 ± 0.31 | 0.0529 ± 0.0036 | |
| Random-fixed | 4.21 ± 0.16 | 4.35 ± 0.28 | 0.0481 ± 0.0039 | |
| Even-stochastic | 2.87 ± 0.07 | 7.20 ± 0.32 | 0.0292 ± 0.0039 | |
| Gamma | 1.40 ± 0.06 | 23.2 ± 1.6 | 0.0384 ± 0.0043 | |
| *full ML decoding for circular feature spaces* | | | | |
| Stochastic | 4.31 ± 0.14 | 5.19 ± 0.25 | 0.0367 ± 0.0041 | |
| Fixed | 6.08 ± 0.88 | 4.54 ± 0.35 | 0.0551 ± 0.0037 | |
| Random-fixed | 4.19 ± 0.17 | 4.40 ± 0.28 | 0.0491 ± 0.0040 | |
| Even-stochastic | 2.87 ± 0.07 | 7.23 ± 0.32 | 0.0292 ± 0.0039 | |
| *excluding swap errors* | | | | |
| Stochastic | 3.57 ± 0.11 | 5.66 ± 0.24 | | |
| Fixed | 2.42 ± 0.06 | 2.97 ± 0.21 | | |
| Random-fixed | 3.61 ± 0.12 | 3.32 ± 0.23 | | |
| Even-stochastic | 2.56 ± 0.06 | 7.43 ± 0.32 | | |
| Gamma | 1.07 ± 0.04 | 31.7 ± 2.1 | | |
| *power law for set size effects* | | | | |
| Stochastic | 7.06 ± 0.39 | 4.64 ± 0.21 | 0.0239 ± 0.0032 | 1.37 ± 0.03 |
| Gamma | 2.33 ± 0.12 | 19.5 ± 1.4 | 0.0258 ± 0.0034 | 1.38 ± 0.03 |

Table S5: Parameter values of ML fits for pooled whole-report data (with a single set of parameters for Experiments 1a & 2a, and for Experiments 1b & 2b).

| Model | $\gamma$ or $K$ | $\omega_1$ | $p_{\mathrm{NT}}$ | $\eta$ |
|---|---|---|---|---|
| *Experiments 1a & 2a* | | | | |
| Stochastic | 3.66 | 4.25 | 0.0204 | |
| Fixed | 3.00 | 2.54 | 0.0631 | |
| Random-fixed | 4.00 | 2.72 | 0.0478 | |
| Even-stochastic | 2.54 | 5.74 | 0.00576 | |
| Gamma | 1.21 | 17.3 | 0.0251 | |
| *Experiments 1b & 2b* | | | | |
| Stochastic | 3.80 | 6.81 | 0.0219 | |
| Fixed | 2.00 | 6.77 | 0.0309 | |
| Random-fixed | 4.00 | 4.79 | 0.0428 | |
| Even-stochastic | 2.64 | 9.20 | 0.0169 | |
| Gamma | 1.09 | 37.3 | 0.0261 | |
| *Experiments 1a & 2a, with random drift* | | | | |
| Stochastic | 3.70 | 4.67 | 0.0181 | 0.0181 |
| Fixed | 3.00 | 2.69 | 0.0614 | 0.0194 |
| Random-fixed | 4.00 | 2.78 | 0.0464 | 0.0114 |
| Even-stochastic | 2.61 | 6.72 | 0.00434 | 0.0232 |
| Gamma | 1.18 | 19.6 | 0.0236 | 0.0151 |
| *Experiments 1b & 2b, with random drift* | | | | |
| Stochastic | 3.99 | 9.62 | 0.0136 | 0.0309 |
| Fixed | 2.00 | 7.60 | 0.0265 | 0.0207 |
| Random-fixed | 4.00 | 5.34 | 0.0314 | 0.0265 |
| Even-stochastic | 2.81 | 14.16 | 0.0100 | 0.0341 |
| Gamma | 0.995 | 67.3 | 0.0196 | 0.0292 |

### References


[1] Ronald van den Berg, Edward Awh, and Wei Ji Ma. "Factorial comparison of working memory models." *Psychological Review* 121.1 (2014), p. 124.

[2] Kirsten CS Adam, Edward K Vogel, and Edward Awh. "Clear evidence for item limits in visual working memory". *Cognitive Psychology* 97 (2017), pp. 79–97.

[3] Paul M Bays. "A signature of neural coding at human perceptual limits". *Journal of Vision* 16.11 (2016), pp. 4–4.

[4] Weiwei Zhang and Steven J. Luck. "Discrete fixed-resolution representations in visual working memory". *Nature* 453.7192 (2008), pp. 233–235.

[5] Ronald van den Berg et al. "Variability in encoding precision accounts for visual short-term memory limitations". *Proceedings of the National Academy of Sciences* 109.22 (2012), pp. 8780–8785.

[6] Daryl Fougnie, Jordan W Suchow, and George A Alvarez. "Variability in the quality of visual working memory". *Nature Communications* 3 (2012), p. 1229.

[7] Robert Taylor and Paul M. Bays. "Theory of neural coding predicts an upper bound on estimates of memory variability". *Psychological Review* Advance Online Publication. http://dx.doi.org/10.1037/rev0000189 (2020).

[8] Paul M Bays. "Noise in neural populations accounts for errors in working memory". *Journal of Neuroscience* 34.10 (2014), pp. 3632–3645.

[9] Alexander S Ecker et al. "Decorrelated neuronal firing in cortical microcircuits". *Science* 327.5965 (2010), pp. 584–587.

[10] Paul M Bays, Raquel FG Catalao, and Masud Husain. "The precision of visual working memory is set by allocation of a shared resource". *Journal of Vision* 9.10 (2009), pp. 7–7.

[11] Sebastian Schneegans and Paul M Bays. "Drift in neural population activity causes working memory to deteriorate over time". *Journal of Neuroscience* (2018), pp. 3440–3417.

[12] Rosanne L Rademaker, Caroline H Tredway, and Frank Tong. "Introspective judgments predict the precision and likelihood of successful maintenance of visual working memory". *Journal of Vision* 12.13 (2012), pp. 21–21.

[13] Paul M Bays et al. "Temporal dynamics of encoding, storage, and reallocation of visual working memory". *Journal of Vision* 11.10 (2011), pp. 6–6.

[14] Paul M Bays, Emma Y Wu, and Masud Husain. "Storage and binding of object features in visual working memory". *Neuropsychologia* 49.6 (2011), pp. 1622–1631.

[15] Nikos Gorgoraptis et al. "Dynamic updating of working memory resources for visual objects". *Journal of Neuroscience* 31.23 (2011), pp. 8502–8511.

[16] Michael S Pratte et al. "Accounting for stimulus-specific variation in precision reveals a discrete capacity limit in visual working memory." *Journal of Experimental Psychology: Human Perception and Performance* 43.1 (2017), p. 6.
